# Perception of Temperature Even in the Absence of Actual Change is Sufficient to Drive Transgenerational Epigenetic Inheritance

**DOI:** 10.1101/2024.12.02.626416

**Authors:** Guy Teichman, Mor Sela, Chee Kiang Ewe, Hadas Gayer, Itai Rieger, Gil Ash, Sarit Anava, Yael Mor, Péter R. Szántó, David H. Meyer, Hila Doron, Or D. Shahar, Vladyslava Pechuk, Hila Gingold, Meital Oren-Suissa, Matthew McGee, Michael Shapira, Björn Schumacher, Oded Rechavi

**Affiliations:** School of Neurobiology, Biochemistry and Biophysics, Wise Faculty of Life Sciences & Sagol School of Neuroscience, Tel Aviv University, Tel Aviv, Israel; Institute for Genome Stability in Aging and Disease, Medical Faculty, University and University Hospital of Cologne, Germany; Cologne Excellence Cluster for Cellular Stress Responses in Ageing-Associated Diseases (CECAD), Center for Molecular Medicine Cologne (CMMC), University and University Hospital of Cologne, Joseph-Stelzmann-Str. 26, 50931 Cologne, Germany; Migal Galilee Research Institute, Upper Galilee, Kiryat Shmona, Israel; Tel-Hal Academic College, Biotechnology Department, Upper Galilee, Kiryat Shmona, Israel; Department of Brain Sciences, Faculty of Biology, Weizmann Institute of Science, Rehovot, Israel; Department of Integrative Biology, University of California, Berkley, California

**Author notes:** These authors contributed equally (Guy Teichman, Mor Sela, Chee Kiang Ewe).

## Abstract

Can processes occurring in one individual’s nervous system influence the physiology of the descendants? Here, we explored the hypothesis that parents’ sensation or perception of environmental cues can influence their offspring, extending across many subsequent generations. We show that in *Caenorhabditis elegans*, temperature perception by the AFD thermosensory neurons initiates a signaling cascade that, directly or indirectly, induces transgenerational changes in RNAi factors, small RNAs, and their target genes. Moreover, we identify secreted factors that enable this neuron-to-germline communication and trace the path of the epigenetic signal. We further model the process mathematically, and the model yields new predictions that we validate experimentally: blocking sensory input dampens RNAi inheritance initiated by exogenous double-stranded RNA (dsRNA). Together, our results demonstrate that sensory perception is sufficient to influence small RNA-mediated heritable gene expression memory.

## Introduction

The nervous system is unrivaled in its ability to gather, process, and integrate information about the environment. Using data collected from sensory and internal organs, neurons orchestrate behavioral and physiological responses and prime future decisions, shaping the organism’s response to environmental cues. In this study, we sought to test the hypothesis that perception – the interpretation of sensory information – not only influences decision-making and physiology within the same generation but also generate long-term, heritable epigenetic memory transmitted across generations. Such a mechanism may be adaptive, although carry-over of aberrant epigenetic information may also be neutral or have negative consequences on health and survival of the animals^1–4^.

*C. elegans* is an exceptional model organism for investigating potential links between the nervous system, the germline, and inheritance. Thanks to the deterministic development of their nervous system, consisting of only 302 neurons, and their mapped neuronal connectome^5–7^, these worms enable studying the nervous system at the circuit and single neuron levels. Moreover, *C. elegans* possess a vast array of small RNA-controlled processes^8,9^. Contrary to the longstanding dogma that DNA being the sole hereditary materials, *C. elegans* can inherit small RNA responses, which may regulate germline gene expression transgenerationally^10,11^.

Small RNA inheritance in *C. elegans* is achieved owing to the active amplification of small RNA responses by the RNA-dependent RNA polymerases (RdRPs) RRF-1 and EGO-1^12–16^. Exogenous and endogenous siRNAs may trigger systemic and heritable gene silencing and a suite of Argonaute proteins regulate the establishment and maintenance of this response^9,12,17–20^. In addition, PIWI protein and their bound piRNAs can induce a transgenerational epigenetic memory of foreign RNA and other germline genes^21,22^.

Environmental stimuli play a role in regulating the synthesis and inheritance of small RNAs. For example, it has been shown that viral^12,23^ and bacterial infections^24^, starvation^25–29^, and changes in temperature^30–33^ can modify the worms’ small RNA pools transgenerationally. In particular, some germline-expressed transgenes, which are generally recognized by the RNAi machinery as foreign genetic elements, undergo accumulative transgenerational epigenetic silencing when worms are grown at 20°C (a non-stressful temperature), but become re-expressed when worms are grown at 25°C (a mildly stressful temperature)^30,32^. Importantly, whether or not the different above-mentioned heritable effects of environmental challenges involve the action of the nervous system is unclear. More specifically, it is unknown whether the effects of temperature on transgene silencing are mediated by specific neuronal thermosensation pathways or by general physiological temperature responses.

Perception occurs following the activation of sensory neurons in response to specific stimuli. The received information is interpreted, inducing changes in behaviors, metabolism, or development that allow the animals to adapt to novel environments. In *C. elegans*, temperature sensation in the non-noxious range is mediated mainly by the AFD pair of sensory neurons, with additional contributions from the AWC, ASI and ASJ neurons^34–40^. Both the AFD and the AWC sensory neurons are postsynaptically connected to the AIY interneurons via chemical synapses^5^, and the AIY, together with other downstream neurons, mediate thermotaxis behavior^35,38,39^. The AFD thermosensory circuit is capable of detecting both temperature changes^41,42^ and long-term ambient temperature^43^, a process requiring the CMK-1 calcium/calmodulin-dependent protein kinase I (CaMKI)^44^. Multiple groundbreaking studies have shown that the AFD thermosensory neurons participate cell non-autonomously in long-term physiological responses to temperatures. AFD activation was shown to be sufficient to induce somatic expression of the transcription factor Heat Shock Factor-1 (HSF-1)^45^, which partakes in the heat-shock response^46^ and promote longevity at 25°C^47–49^.

Previous studies suggest that certain neuronal processes initiated in parents could continue to affect their F1 progeny^50–52^. For example, parental exposure to pheromones was shown to affect the children’s reproduction and developmental rates^52^. However, these effects did not last beyond the F1 generation – an exemplar of short-term *intergenerational* inheritance, as opposed to perduring *transgenerational* inheritance that affects descendants not directly exposed to the original trigger (for example *in utero* exposure)^53^. In contrast, other studies have demonstrated that small RNAs synthesized in neurons can affect gene expression and behavior transgenerationally (≥ 3 generations) in *C. elegans*^54^. Moreover, it has been shown that neuronal hairpin-derived dsRNAs can be transmitted to the germline and induce transgenerational gene silencing^55^. It is crucial to note that these studies relied on transgenic manipulation of the small RNA pathway in neurons; a direct causal relationship between brain activity or sensory perception and transgenerational small RNA inheritance has yet to be established.

In this work, we used multiple independent experimental systems, coupled with computational modeling, to separate the direct biophysical effects of ambient temperature from its neuronal perception. Our findings reveal that the neuronal perception of temperature via AFD neurons, directly or indirectly, influences the expression of small RNA machinery in the germline, affecting transgenerational epigenetic inheritance. This neuronal regulation of small RNA machinery depends on CMK-1/ CaMKI, the neuropeptide FLP-6, serotonergic signaling, and the transcription factor HSF-1. Our findings imply that sensory perception regulates transgenerational inheritance of gene expression memory, which may influence the animal’s survival fitness in changing environments.

## Results

### Disrupting thermosensation affects transgenerational gene silencing

To study whether temperature perception can affect epigenetic inheritance, we first investigated the effect of AFD-mediated temperature perception on temperature-dependent germline transgene silencing. We used worms that carry a low-copy integrated *gfp* transgene under the control of a germline promoter (*bnIs1[pie-1p::gfp::pgl-1+unc-119*(+)]), which were previously described^56^ (see **Figure 1A** and **Materials and Methods**). This transgene is constitutively silenced at 20°C. However, we observed that growing the worms at elevated temperature (25°C) for at least 24 hours rapidly and robustly triggered *gfp* expression from the transgene (**Figure 1B; Extended Figure 1**). This upregulation of *gfp* is accompanied by downregulation of siRNAs targeting the transgene at high temperature (see below). Shifting the worms back to low temperatures (20°C) resulted in a gradual, cumulative silencing response over 3-7 generations (**Figure 1C**). The expression of *bnIs1* was de-repressed in mutants lacking *hrde-1*, a gene encoding a germline nuclear Argonaute involved in RNAi inheritance^17^, when raised at 20°C (**Figure 1D**). This is consistent with previous study that the germline nuclear RNAi pathway normally represses heat-induced transgenerational inheritance^33^. Hence, the *bnIs1* transgene serves as a sensitive, binary reporter for dissecting the genetic pathway that governs temperature-dependent, heritable gene regulation mediated by RNAi. As a complementary readout, we examined the expression of an endogenous gene *scrm-4*, encoding phospholipid scramblase, which we found to be heat-inducible and regulated by siRNAs, like *bnIs1* (**Figure 1E; Extended Figure 2A**). Importantly, *scrm-4* also exhibited transgenerational accumulation of silencing when transitioning from 25°C to 20°C (**Extended Figure 2B**), and its expression was de-repressed in an *hrde-1* mutant background (**Figure 1F**). Concordantly, siRNAs targeting *scrm-4* is significant enriched in HRDE-1 complexes (2.47 fold enrichment relative to co-IP input controls) and these siRNAs is depleted in *hrde-1(tm1200)* mutants (log2FC = -1.60; q < 0.0001)^9^.

**Figure 1:**
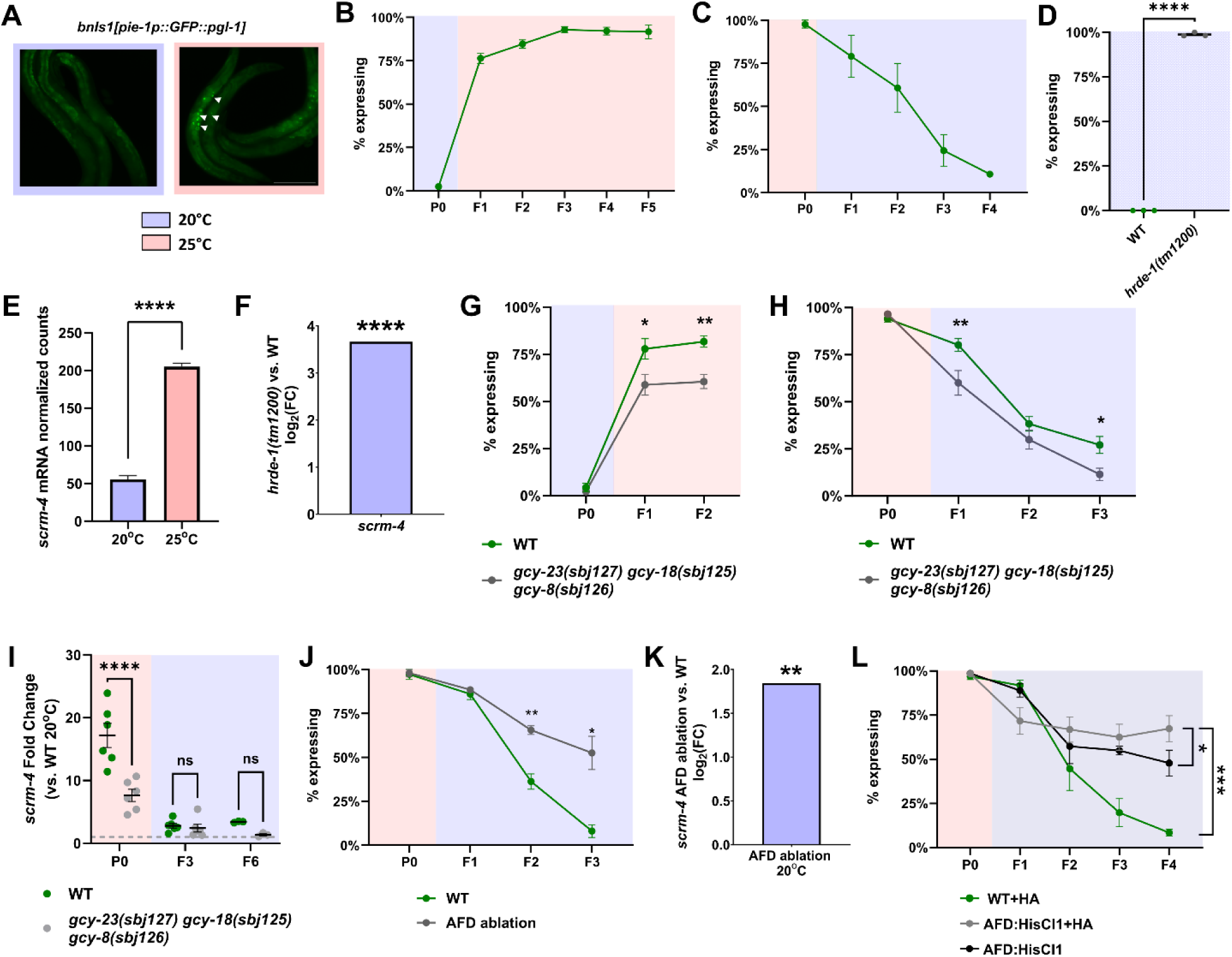
The AFD thermosensory neurons control temperature-dependent small RNA-mediated gene silencing. (A) Representative images of worms containing the *bnIs1* transgene, grown in either high (25°C) or low (20°C) temperatures. (B, C) The archetypal dynamics of transgene silencing and de-silencing upon transition between temperatures. (D) Knocking out *hrde-1* de-represses *bnIs1* transgene (E) *scrm-4* is upregulated at 25°C compared with 20°C. (F) Knocking out *hrde-1* de-represses *scrm-4* (G, H) Knocking out AFD-specific guanylyl cyclases *gcy-8/18/23* promotes transgene silencing upon transition between temperatures. (I) Knocking out AFD-specific guanylyl cyclases *gcy-8/18/23* prevents de-repression of scrm-4. (J) AFD ablation suppresses transgene silencing upon transition from 25°C to 20°C. (K) AFD ablation causes upregulation of *scrm-4* at 20°C. (J) Hyperpolarization of the AFD neuron suppresses transgene silencing upon transition from 25°C to 20°C. In all relevant panels, error bars indicate mean ± SEM. Statistical significance was determined using mixed effect model followed by post-hoc linear contrast with two-stage FDR correction. (*) indicates q < 0.05, (**) indicates q < 0.01, (***) indicates q < 0.001, and (****) indicates q < 0.0001 (see **Materials and Methods**).

We examined mutants carrying null mutations in the three genes encoding AFD-specific guanylyl cyclases *gcy-8*, *gcy-18*, and *gcy-23* (henceforth referred to as “AFD triple mutants” for brevity). These partially redundant guanylyl cyclases are expressed *exclusively* in the AFD neurons^57–59;^ knocking out all three factors completely eliminates thermosensation in the AFD neurons and the mutant animals are atactic on thermal gradients^60–62^. We note that, during the course of this study, we tested multiple strains carrying the triple mutations. We found that the strain PY9248 (*gcy-23(oy150) gcy-8(oy44) gcy-18(nj38)*) may carry background mutation(s), causing a mutator phenotype with enhanced transposon mobilization, potentially regulated by small RNA^63^. As such, we generated a new triple AFD mutant carrying identical loss-of-function alleles using CRISPR/Cas9. The resulting strain, BJS1456, was backcrossed to the N2 reference strain twice to remove off-target mutations. Using this newly generated strain, we found that AFD triple mutants failed to fully de-repress the *bnIs1* transgene during the transition from 20°C to 25°C (**Figure 1G**). Moreover, when transitioning from 25°C to 20°C, AFD triple mutants rapidly silence the transgene over 2-3 generations – a much shorter epigenetic memory compared to wild type (**Figure 1H**). We observed a similar phenotype by measuring *scrm-4* expression by RT-qPCR: *scrm-4* was less strongly induced at 25°C in AFD mutants than in wild-type animals (**Figure 1I**). Moreover, we found that *scrm-4* is targeted by piRNAs in the germline (target sites were predicted using the default “stringent” setting from piRTarBase) (**Extended Figure 2C**), and that loss of PRG-1/PIWI leads to upregulation of *scrm-4* at 20°C; this effect is further exacerbated at 25°C (**Extended Figure 2D**). Remarkably, this is completely rescued in the AFD triple mutant background (**Extended Figure 2D)**, likely mediated by downstream WAGOs, which are known to mediate aberrant gene silencing when the piRNA pathway is disrupted^64^. Together, these results suggest that loss of temperature sensation in the AFD neurons in particular enhances small RNA-mediated gene silencing and influences transgenerational epigenetic inheritance. Our results do not preclude the possibility that disruption of AFD neurons may trigger a systemic stress response that indirectly affects RNAi pathways.

To substantiate our conclusion, we genetically ablated AFD neurons by expressing a reconstituted caspase under the control of AFD-specific *gcy-8* promotor^39^ (**Figure 1J**). We found that worms lacking AFD neurons showed delayed *bnIs1* silencing when transitioning from 25°C to 20°C. Moreover, AFD ablation upregulates *scrm-4* expression (**Figure 1K**). The phenotypic differences between AFD-ablated animals and AFD triple mutants suggest that additional mechanisms within AFD likely contribute to the regulation of germline gene expression. In wild-type animals, intracellular calcium levels are inversely correlated with growth temperatures^43^. It was reported that AFD triple mutants exhibited a higher baseline level of intracellular calcium^65^, assimilating animals grown at low temperature. This chronic high-calcium state that fails to change in response to temperature switch in triple *gcy* mutants, in contrast to the wholesale loss of AFD neurons, may explain the distinct phenotypes we observed.

To further examine the impact of AFD neuronal activity on temperature-dependent transgene expression, we utilized the *Drosophila* histamine-gated chloride channel HisCl1 to reversibly hyperpolarize the AFD neurons^66,67^. We generated three independent lines expressing HisCl1 under the control of AFD-specific *gcy-8* promoter (see **Materials and Methods**). Worms carrying AFD:HisCl1 experience hyperpolarization of the AFD neurons when exposed to exogenous histamine (**Extended Figure 3A, B**). We then examined the impact of chronic HisCl1-driven AFD silencing, maintained through histamine supplementation, on temperature-dependent small RNA-mediated transgene silencing. We found that AFD:HisCl1 worms grown on histamine exhibited delayed silencing dynamics of the GFP reporter, similar to AFD ablation (**Figure 1L; Extended Figure 3C)**. While the wild-type control groups exhibited the expected progressive silencing of the GFP reporter over the course of 3-6 generations, the experimental group maintained stable GFP expression in the population for at least 7 generations (**Extended Figure 3C)**. Moreover, we found that histamine treatment of AFD:HisCl1 worms for one generation was sufficient to delay transgene silencing (**Extended Figure 3D, E;** see purple group). Importantly, expressing HisCl1 under the control of *sra-7* promotor in ASK neurons^68^, which are not involved in temperature sensation (see **Extended Table 1**), has no effect on germline gene silencing, regardless of the presence of exogenous histamine (**Extended Figure 3F**), suggesting that our observed effects are specific to the AFD neurons.

It is noteworthy that worms carrying multicopy AFD:HisCl1 extrachromosomal array, even in the absence of histamine, exhibited an intermediate effect between the experimental group and the wild-type control groups. This may be a result of “leakiness” of the HisCl1 channels in the absence of histamine, which has been previously reported^69,70^. In contrast, expressing the channel from a single copy transgene (SC-AFD:HisCl1, see **Materials and Methods**) does not affect transgene silencing (**Extended Figure 3G),** likely due to insufficient channel expression to induce substantial neuronal hyperpolarization. Notably, all prior studies reporting effective HisCl1-mediated neuronal silencing employed high-copy-number expression systems^66,67,69–71^. Nevertheless, we leverage the leakiness of the multicopy AFD:HisCl1 array to investigate long-term effects of transient AFD disruption. We found that AFD:HisCl1 induces changes in temperature-dependent small RNA-mediated transgene silencing that lasts for at least 3 generations after the transgene is lost (**Extended Figure 3H, I).** Taken together, these results imply that short term (1 generations) manipulation of activity in the AFD neurons can cause long-term (> 3 generations) transgenerational expression memory mediated by small RNAs in the germline.

### Thermosensation controls germline small RNA-mediated gene silencing via neuropeptide signaling

To understand the mechanisms through which temperature perception affects gene silencing in the germline and epigenetic inheritance, we examined additional mutants that lack components downstream to GCY-8/18/23 (**Figure 2A**). We started by inspecting mutants defective in the gene *cmk-1*/*CaMKI*. CMK-1 is a conserved calcium/calmodulin-dependent kinase expressed primarily in sensory neurons, and most prominently in the AFD neurons. In fact, compared to other neuronal classes, AFD exhibits both the highest *cmk-1* level and constitutes the largest fraction of isolated cells that express *cmk-1* based on the CeNGEN single cell sequencing dataset (**Extended Figure 4A**)^58^. CMK-1 is required for long-term temperature adaptation in the AFD neurons^43,44,72^. We found that mutants of *cmk-1* exhibited a delayed transgenerational silencing response when transitioning from 25°C to 20°C, i.e., prolonged heat-induced expression memory, similar to AFD ablation and AFD:HisCl1 (**Figure 2B**).

**Figure 2:**
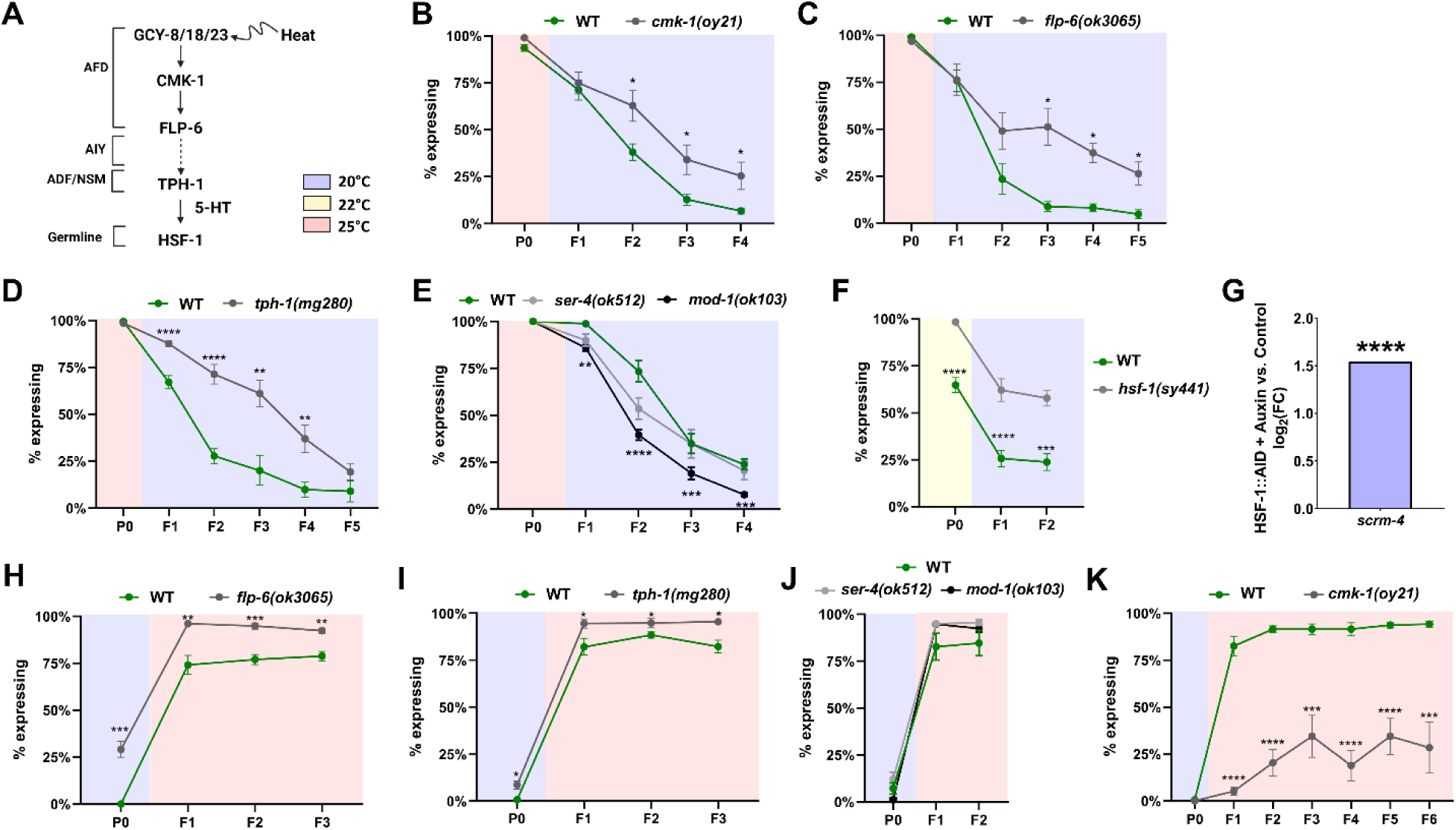
The thermosensory circuit and HSF-1 regulates temperature-dependent gene silencing and transgenerational epigenetic inheritance. (A) A scheme of the AFD thermosensory circuit and its downstream components. (B-D) Knockout of components in the AFD thermosensory circuit inhibits transgene silencing upon transition to a low temperature. (E) Knocking out *mod-1*, but not *ser-4*, promotes transgene silencing upon transition to low temperature. (F) Knocking out *hsf-1* inhibits transgene silencing upon transition to a low temperature. (G) Germline-specific degradation of HSF-1 causes upregulation of scrm-4. (H-K) Knocking out of *ser-4*, *mod-1*, or other components in the AFD thermosensory circuit, with the exception of *cmk-1* (H), does not abolish the archetypal transgene de-silencing upon transition to a high temperature. In all relevant panels, error bars indicate mean ± SEM. (*) indicates q < 0.05, (**) indicates q < 0.01, (***) indicates q < 0.001, and (****) indicates q < 0.0001 (see **Materials and Methods**).

Previous studies have shown that CMK-1 can phosphorylate the CREB homolog CRH-1 at the serine 48 residue^47,73^. Moreover, it was demonstrated that CMK-1 phosphorylates CRH-1 in the AFD neurons in a temperature-dependent manner, promoting the expression of the neuropeptide-encoding gene *flp-6*^47,74^. FLP-6 subsequently mediates communication between the AFD thermosensory neurons and the AIY interneurons^47^. We therefore hypothesized that the AFD-AIY thermosensory circuit could regulate temperature-dependent transgene expression in the germline. When examining mutants of *flp-6*, we found that they exhibited an extended transgenerational expression memory when transitioning from high to low ambient temperatures, as observed in *cmk-1(-)* (**Figure 2C**). This phenotype persisted for at least 10 generations following the transition to 20°C (**Extended Figure 4B**).

Taken together, our results suggest a model whereby CMK-1, functioning downstream of GCY-8/18/23 in AFD, activates neuropeptide FLP-6, which leads to transgenerational silencing of germline transgene; however, we cannot exclude the involvement of other sensory neurons outside of AFD. While our results do not preclude the possibility that disrupting neuronal signaling triggers a systemic physiological response that ultimately influences the RNAi machinery in the germline, they nevertheless provide strong evidence that regulatory information originating in sensory neurons can induce perduring gene expression changes that persist across generations.

### Change in ambient temperatures and HSF-1 activity affect small RNA inheritance in both sensation-dependent and -independent manners

HSF-1, a conserved transcription factor involved in development and stress response, has previously been shown to determine the “epigenetic inheritance state” of worms by regulating small RNA factors in the germline. In particular, low activity of HSF-1 reduces small RNA-dependent silencing of transgenes and RNAi inheritance, while HSF-1 over-expression supports small RNA-mediated gene silencing^75^. Notably, the AFD neurons have been shown to regulate germline activation of HSF-1 cell non-autonomously through serotonergic signaling mediated by serotonin receptor SER-1^46,49^. We therefore hypothesized that HSF-1 may similarly regulate the effects of neuronal sensation on small RNA inheritance.

To investigate the involvement of serotonin release in small RNA-mediated gene silencing, we examined whether mutants of the tryptophan hydroxylase *tph-1* or the serotonin receptor *ser-1* show modified temperature-dependent transgene expression. We found that *tph-1(-)* mutants exhibited a strong delayed transgene silencing (**Figure 2D**), persisting for at least 10 generations following the transition to 20°C (**Extended Figure 4C**). However, we could not detect any difference in transgene expression in *ser-1(-)* mutants (**Extended Figure 4D**). We next tested the involvement of two additional serotonin receptors – MOD-1 and SER-4 – in this process. Upon the shift from 25°C to 20°C, *ser-4(-)* mutants showed no effect, however, surprisingly, *mod-1(-)* mutants show a rapid loss of transgenerational memory – a phenotype opposite to that of *tph-1(-)* mutant (**Figure 2E**). These findings suggest that the effect observed in *tph-1(-)* mutants may involve additional serotonin receptors not examined in this study, and that serotonin receptors may act in combination to regulate germline gene expression. Supporting this notion, many previous studies have identified opposing functions of different serotonin receptors^76–78^.

Next, to investigate the involvement of HSF-1, we examined the temperature-dependent silencing dynamics of *hsf-1* partial loss-of-function mutants (*hsf-1* null mutants are lethal^79^). Since *hsf-1* hypomorph mutants (*sy441*) are hypersensitive to heat stress (they die at 25°C), we conducted these experiments within a narrower temperature range (20-22°C). We found that *hsf-1(sy441)* mutants display a reduced gene-silencing phenotype reminiscent of *flp-6(-)* and *tph-1(-)* mutants (**Figure 2F**). Moreover, we found that degrading HSF-1 specifically in the germline using the Auxin-inducible degron (AID) system led to de-silencing of *scrm-4* (**Figure 2G**). While perturbing AFD neurons and downstream serotonergic and HSF-1 signaling may trigger a systemic stress response that indirectly modulates RNAi pathways, our results nevertheless indicate that heritable transgene silencing in the germline, directly or indirectly, depends on serotonin release, likely from ADF and NSM downstream of AFD in the thermosensory circuit, as well as HSF-1 activity in the germline (more below).

### Mutants of *cmk-1* exhibit a high-variance gene-silencing response at high ambient temperature

Intriguingly, the different mutants we examined showed distinct silencing dynamics. When we transitioned the worms from low temperature (20°C) to high temperature (25°C), we did not detect obvious difference between wild-type worms and *tph-1(-)*, *mod-1(-), ser-4(-),* and *flp-6(-)* (**Figure 2H-J**), with all strains achieving near-complete de-silencing of the GFP reporter within 2-3 generations (note that *tph-1(-)* and *flp-6(-)* mutants started from a higher baseline). In contrast, AFD triple and *cmk-1(-)* mutants showed enhanced silencing of *bsIn1* transgene at 25°C (**Figure 1G** and **Figure 2K**).

Focusing on *cmk-1(-)* mutants, we observed an unusual high-variance response when transitioning from low to high temperature: some worm lineages exhibited rapid de-silencing of the GFP reporter, others maintained complete silencing for at least 7 generations, and still others repeatedly fluctuated between the two extremes (**Figure 2K, Extended Figure 5A, B**). We also observed downregulation of *scrm-4* in *cmk-1(-)* grown at 25°C (**Extended Figure 5C**). These results suggest that different factors, perhaps in neurons outside of AFD, coordinate the responses to changes in temperature, depending on whether the worms transition up or down the temperature gradient.

### Characterizing the effects of neuronal temperature sensation on the mRNA and small RNA expression

Our foregoing results demonstrate that the AFD thermosensory circuit has a major impact on transgene silencing and epigenetic inheritance at different temperatures. To understand the underlying mechanism, we examined gene expression in AFD-ablated worms and found that epigenetic regulators tended to be downregulated in the mutants compared to wild type (**Figure 3A**). Moreover, targets of MUT-16, which is required for small RNA amplification, tended to be upregulated in AFD-ablated animals (**Figure 3B**), suggesting defects in the small RNA pathways, which cause de-silencing of small RNA targets.

**Figure 3:**
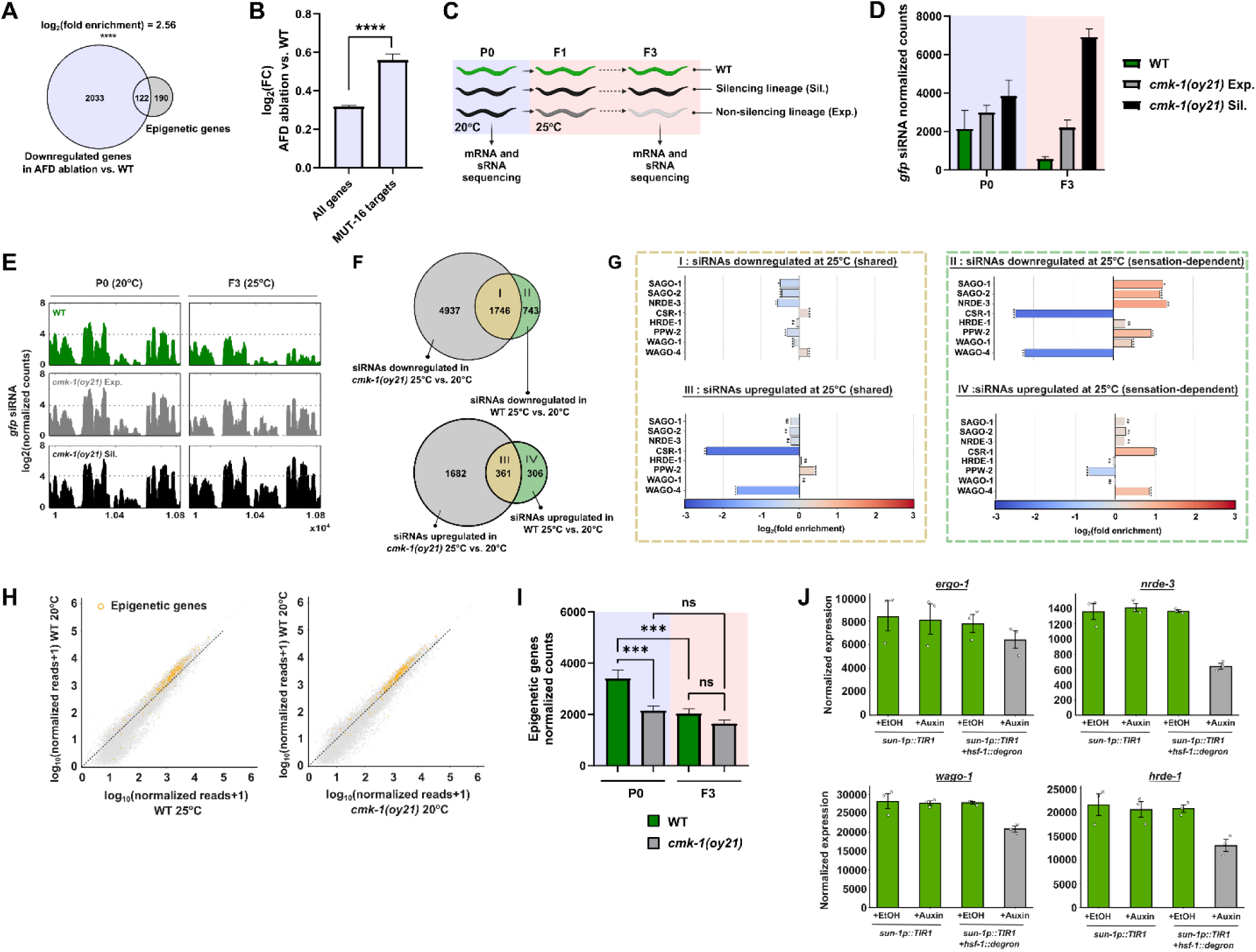
Temperature perception regulates epigenetic gene expression and small RNA pathways. (A, B) AFD ablation causes down regulation of epigenetic genes and upregulation of MUT-16 targets. (C) Experimental scheme. We sequenced mRNA and small RNAs from wild-type and *cmk-1* mutant worms across generations and temperatures. We grew worms at 20°C for multiple generations and then transferred them to 25°C for three generations. We established independent lineages from P0, tracking GFP expression throughout. We sequenced worms at P0 (20°C) and F3 (after three generations at 25°C). For F3 *cmk-1(-)* mutants, we selected “expressing” (highest GFP expression) and “silencing” (lowest GFP expression) lineages, based on their F2 phenotypes. This allows comparison of differential responses to temperature change. (D, E) Anti-*gfp* siRNA levels are downregulated upon transition to 25°C in a CMK-1-dependent manner. Reads were normalized using the Relative Log Expression method. Silencing lineages of *cmk-1(-)* showed higher levels of anti-*gfp* siRNA than expressing lineages. (F) Growth at 25°C induces changes in small RNA pools in both a CMK-1/sensation-dependent and -independent manner. Shown are Venn diagrams of the siRNAs significantly up-or down-regulated in 25°C in wild-type worms only (green), *cmk-1(-)* mutants only (grey), or both (yellow). (G) CMK-1/sensation-dependent changes in siRNAs following temperature transition are enriched for specific Argonaute targets. Shown are fold enrichment and depletion values for downregulated (bottom) or upregulated siRNAs (top), and siRNAs that change in a sensation-dependent (right half) or -independent manner (left half). Roman numerals (I-IV) correspond to (H, I) The expression of epigenetic-related genes is increased in worms grown at 20°C (Y-axis) compared to worms grown in 25°C (X-axis). Epigenetics genes are downregulated in *cmk-1(-)* compared with wild type. In (H), each dot represents a protein-coding gene. Orange dots represent epigenetic-related genes. Dotted line represents the diagonal (where expression values equal between the two conditions). Reads were normalized using the Relative Log Expression method. (J) The expression of small RNA factors is decreased upon germline-specific degradation of HSF-1. Shown are the Relative Log Expression-normalized expression levels (geometric mean ± GSEM) of two germline-specific Argonaute genes (*hrde-1* and *wago-1*) and two ubiquitous Argonautes (*ergo-1* and *nrde-3*). Each dot corresponds to a single biological replicate (*) indicates q < 0.05, (**) indicates q < 0.01, (***) indicates q < 0.001, and (****) indicates q < 0.0001 (see **Materials and Methods**).

Next, given the unusual gene silencing behaviors of *cmk-1(-)* mutants compared to other sensory mutants we analyzed, we sought to characterize the high-variance response that we observed in *cmk-1(-)* mutants upon transition from low to high temperatures (**Figure 2K** and **Extended Figure 5A, B**). To that end, we sequenced small RNAs and mRNAs from wild-type and *cmk-1(-)* worms before and after transitioning from a low temperature (20°C) to a high temperature (25°C). For the mutant worms, we isolated both “expressing” lineages (i.e., lineages in which GFP silencing was abolished at high temperature) and “silencing” lineages (i.e., lineages in which GFP silencing was maintained at high temperature) (**Figure 3C** and **Materials and Methods**).

Interestingly, in the small RNA data sequenced from the P0 generation (20°C), we did not detect changes in anti-*gfp* siRNA levels between “expressing” and “silencing” lineages (log2FC = -0.11, q = 0.99). In F3 (25°C), however, we detected a significant difference in anti-*gfp* siRNAs (**Figure 3D, E; q < 0.006**), as well as changes in siRNAs known to be associated with specific Argonautes (**Extended Tables 2 and 3**). Despite changes in the siRNA pools, we did not detect any significant differences in mRNA expression between the GFP-silencing and GFP-expressing lineages. We did not observe any changes in anti-*pgl-1* siRNA or *pgl-1* mRNA levels. These results indicate that (1) the bimodal silencing phenotypes observed in *cmk-1(-)* mutants emerge only after the transition to 25°C and cannot be directly predicted from the small RNA pools or gene expression profiles of the P0 ancestors, and (2) the *gfp* transgene appears to be stochastically silenced independent of gene expression states of other loci.

In agreement with the reduction in the gene-silencing capacity we observed in *cmk-1(-)* mutants, we detected a significant reduction in the fraction of siRNAs targeting protein-coding genes in *cmk-1(-)* mutants grown in both 20°C and 25°C (**Extended Figure 6A).** We found siRNA changes that are both sensation-dependent (i.e., occur in wild-type worms but not in *cmk-1(-)*) and -independent (i.e., common to both wild-type worms and *cmk-1(-)*) (**Figure 3F**). While the siRNAs changing in a sensation-independent manner were not enriched for targets of particular Argonaute proteins, we found that siRNAs downregulated in a sensation-dependent manner were enriched for small RNAs bound to the ubiquitous Argonautes SAGO-1, SAGO-2, and NRDE-3, as well as the germline-specific Argonautes PPW-2 and WAGO-1 (**Figure 3G**)^9^. Moreover, we found that siRNAs upregulated in a sensation-dependent manner were enriched for CSR-1- and WAGO-4-bound small RNAs (**Figure 3G**), which were previously shown to target a similar cohort of constitutively-expressed germline genes^9,80,81^. Of particular interest, WAGO-4 has previously been found to mediate transgenerational epigenetic inheritance^81,82^. Additionally, CSR-1 is thought to protect germline transcripts from piRNA silencing^80,83–85^, which may be enhanced in *cmk-1(-)* (more below). Together, these results indicate that, while high temperatures can non-specifically disrupt small RNA-mediated gene silencing, the neuronal perception of ambient temperature may regulate specific small RNA pathways in the germline.

We then proceeded to examine changes in gene expression in response to changes in ambient temperature and loss of *cmk-1*. As expected, Loss of *cmk-1* leads to misregulation of genes involved in sensory non-motile cilium assembly (GO:1905515; 15 observed vs. 3.9 expected), as well as genes associated with transcriptional and translational regulation (**Extended Figure 6B, C**). By correlating siRNA and mRNA expression, we did not detect a clear genome-wide relationship across the different samples, beyond specific examples such as foreign genetic elements (e.g., *bnIs1*) and *scrm-4* (**Extended Figure 6D**), highlighting the complexity of small RNA pathways in *C. elegans*^86^.

In agreement with previous studies^33^, we found that a large group of epigenetic factors, comprising multiple Argonaute genes (including the germline-specific Argonaute genes *hrde-1*, *wago-1*, *wago-4*, and *ppw-1*), as well as multiple RNA-dependent RNA polymerase genes (*rrf-1, rrf-3,* and *ego-1*), are significantly downregulated at high temperatures in wild-type animals (**Figure 3H**). Crucially, these epigenetic genes, many of which are involved in post-transcriptional gene silencing (GO:0035194), are chronically downregulated in *cmk-1(-)* mutants regardless of growth temperature (**Figure 3H, I, Extended Figure 6C, E,** and **Extended Table 4**). Finally, we found that many of these small RNA factors (75 out of 312 epigenetic genes) are similarly downregulated following germline depletion of HSF-1 (**Figure 3J** and **Extended Table 4**)^87^, further supporting the involvement of germline HSF-1 in the small RNA response to temperature sensation.

### Temperature sensation affects the piRNA pathway and innate immunity

While analyzing small RNA sequencing data from *cmk-1(-)* mutants, we noted that, unlike wild-type animals, which showed downregulation of 21U piRNAs at high temperature^32^, *cmk-1(-)* mutants exhibited increased piRNA abundance upon transition from 20°C to 25°C (**Figure 4A**). Concordantly, we observed downregulation of piRNA-dependent 22G-RNA protein-coding targets in *cmk-1(-)* mutants (**Figure 4B**). Because CSR-1 normally protects germline genes from piRNA-mediated silencing^85,88^, we next examined CSR-1 protein-coding targets and found that they also tended to be downregulated in *cmk-1(-)* mutants; this effect was observed at both 20°C and 25°C, despite no drastic change in 21U piRNA abundance at 20°C (**Figure 4C**). Loss of PRG-1 leads to the aberrant silencing of histone genes by siRNAs whose biogenesis depends on CSR-1^64^. In *cmk-1(-)* mutants, we found that these anti-histone siRNAs are downregulated, consistent with enhanced piRNA pathway (**Figure 4D**).

**Figure 4:**
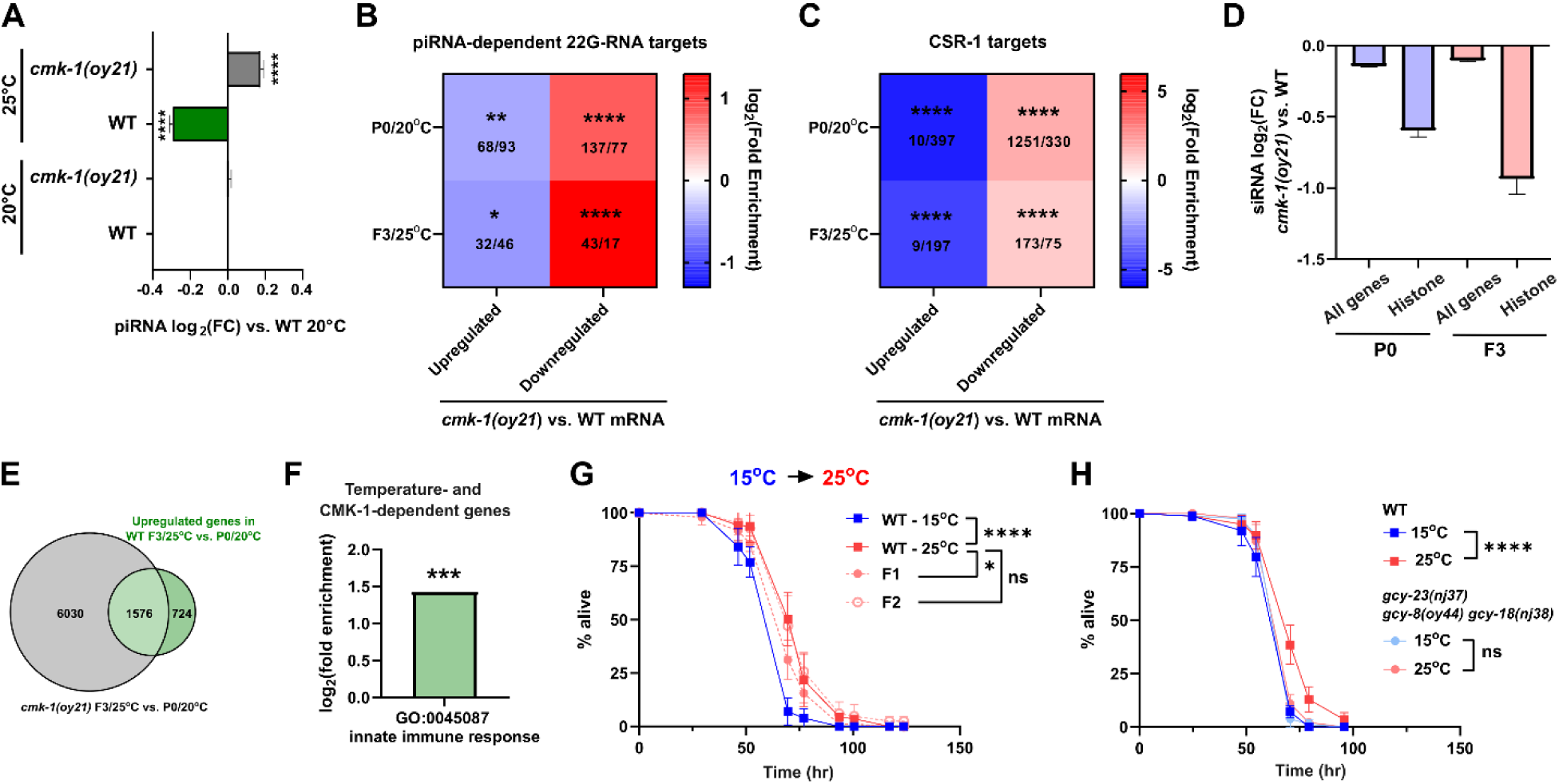
CMK-1 and thermosensation affect piRNA pathway and innate immunity. (A) piRNAs are downregulated at 25°C in wild-type worms but are upregulated at high temperatures in *cmk-1(-)* mutants. Shown are the log2(FC) values of individual piRNAs in comparison to wild-type worms at 20°C. (B, C) piRNA-dependent 22G-RNA and CSR-1 targets are downregulated in *cmk-1(-)* in 20oC and 25°C. (D) siRNA targeting histones are downregulated in *cmk-1(-)* in 20oC and 25°C. (E, F) Gene Ontology enrichment analysis of temperature- and sensory-dependent genes (n = 735) reveals significant enrichment of immune-related genes. (***) indicates q < 0.001. (G) PA14 resistance of wild-types worms transferred from 15°C to 25°C. Cultivation temperature causes integrational effect on innate immune response (H) PA14 resistance of wild type and *gcy-8/18/23* triple mutants at 15°C and 25°C. For all relevant panels, error bars indicate mean ± SEM. In (G) and (H), (*) indicates p < 0.05 and (****) indicates p < 0.0001 by log-rank test.

Previous study demonstrated that infection with pathogenic bacteria suppresses downregulation of the piRNA pathway at high temperature^32^. In addition, loss of CSR-1 leads to increased resistance to bacterial infection^80^. By performing GO enrichment analysis, we found that temperature- and sensation-dependent genes (i.e., genes upregulated in wild-type worms but not in *cmk-1(-)* mutants upon the transition from 20°C to 25°C) are enriched for innate immune functions (**Figure 4E, F**). We therefore hypothesized that temperature sensation via AFD may influence the animals’ susceptibility to infection, potentially by modulating the piRNA/CSR-1 regulatory axis. It has been previously demonstrated that high temperature activates innate immunity by PMK-1/p38 MAP kinase^89,90^; consistently, we found that wild-type worms reared at 25°C (> 5 generations) were more resistant to *Pseudomonas aeruginosa* (PA14) infection compared to those reared at 15°C (**Figure 4G**). Note that the survival assays were performed on the full lawn of PA14, as previous described^91^, to rule out differences in pathogen sensing and avoidance. Interestingly, the progeny of parents grown at 15°C showed a reduced resistance at 25°C compared to worms continuously maintained at 25°C for more than five generations (**Figure 4G**); however, this effect does not appear to persist beyond F1 – an intergenerational effect (**Figure 4G**). In contrast to wild-type animals, triple *gcy* mutants lacking temperature perception showed no temperature-dependent differences in infection susceptibility, instead behaving similarly to animals maintained at 15°C. (**Figure 4H**), indicating a role of thermosensation in promoting immunity at high temperature.

In summary, while exposure to high temperature appears to disrupt worms’ small RNA pools in a non-specific manner, loss of CMK-1-mediated sensory signaling leads to targeted changes in the composition of small RNAs. Of note, AFD ablation leads to downregulation of epigenetic factors, which may subsequently impair germline siRNA-mediated gene silencing. Additionally, we found that intact AFD thermosensation is important for activating innate immunity against pathogenic bacteria. The overlapping transcriptional signatures observed in *cmk-1(-)* and AFD-ablated worms suggest that AFD-mediated temperature sensation elicits specific epigenetic responses in the germline, affecting the expression of many endogenous targets beyond transgene. That said, we cannot rule out additional contributions from other neurons (see below).

### Predicting the impact of changes in ambient temperatures and sensory input on small RNA-mediated gene silencing

Next, we turned to a mathematical modeling-based approach to further understand how neuronal sensation influences different components of the epigenetic system, resulting in different gene-silencing kinetics. Multiple theoretical models have been proposed to explain the dynamics of transgenerational inheritance of epigenetic information^75,92–94^. To further explore the impact of temperature and temperature perception on transgenerational inheritance, we expand on the recently published mathematical model developed by Karin et al.^93^, based on well characterized functional properties of siRNA and H3K9me3 (**Figure 5A**, **Extended file 1** and **Materials and Methods** for the detailed assumptions and parameters of the model).

**Figure 5:**
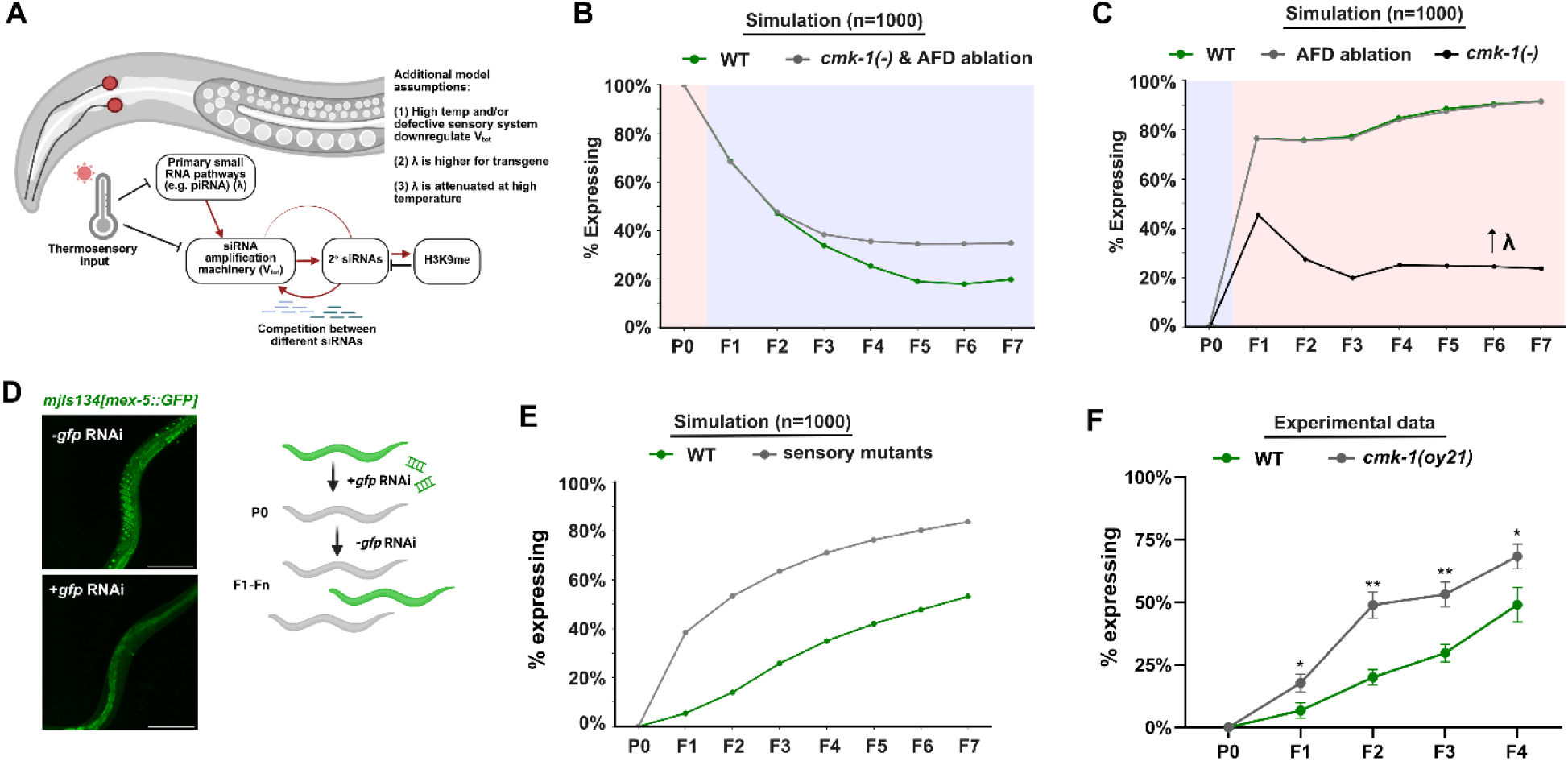
Theoretical model of sensory-dependent small RNA-mediated gene silencing predicts heritable response to exogenous dsRNA. (A) The scheme and added assumptions of the expanded mathematical model describing the impacts of temperature perception on transgenerational small RNA inheritance. The basic model was previously described. (B) The simulated dynamics of transgene silencing after transition from high to low temperatures, in wild-type worms (green) and sensory mutants (grey). (C) The simulated dynamics of transgene de-silencing after transition from low to high temperatures in wild-type worms (green), AFD ablation (grey), and *cmk-1(-)* (black). (D) RNAi inheritance assay based on silencing *mjIs134[mex-5::gfp]* transgene by exogenous dsRNA targeting *gfp*. Scale bar = 200 µm. (E) The simulated dynamics of RNAi inheritance in wild-type worms (green) and temperature sensory mutants (grey). (F) Disrupting sensory perception by knocking out cmk-1 is sufficient to dampen transgenerational inheritance of RNAi responses initiated by exogenous dsRNA. Error bars indicate mean ± SEM. (*) indicates q < 0.05, (**) indicates q < 0.01, (***) indicates q < 0.001, and (****) indicates q < 0.0001 (see **Materials and Methods**). In (B, C), the graphs indicate the mean percentages of worms positive for GFP expression for each generation and experimental condition (number of simulations = 1000).

Our revised model makes three additional assumptions based on our RNA sequencing data: (1) high temperatures (or defective thermosensation) reduce the silencing capacity of the worms; (2) the *gfp* reporter gene is targeted by stochastic silencing events at a high frequency; and (3) the frequency of stochastic silencing events against *gfp* is reduced by high temperatures (or *cmk-1* loss-of-function).

Our simulations show that the model predicts the empirically observed kinetics of *gfp* silencing following temperature transitions: a stochastic and cumulative silencing response upon transition from high to low temperature (**Figure 5B**) and a rapid, uniform de-silencing response upon transition from low to high temperature (**Figure 5C**). Moreover, in agreement with our RNA sequencing results, our simulations suggest that a perception-dependent reduction in silencing capacity is sufficient to explain the reduced transgene-silencing phenotype we observed in multiple temperature sensory mutants, including AFD ablated animals, *cmk-1(-)*, *flp-6(-)*, and *tph-1* mutants. (see **Figure 5B, C** for simulated results; see also **Figure 2** for experimental results). Crucially, increasing piRNA-mediated silencing, represented by λ, recapitulated high-variance gene-silencing response observed in *cmk-1(-)* at 25°C (**Figure 5C**). The broad expression of cmk-1 in the nervous system and the unusual silencing dynamics of the mutants prompt us to test its role in AFD neurons. We expressed *cmk-1* under the control of AFD-specific *gcy-8* promotor in *cmk-1(-)* background and found that the rescuing strain showed a stronger silencing phenotype than the mutant alone (**Extended Figure 7A, B**), suggesting that CMK-1 in AFD promotes silencing, while CMK-1 in other neurons suppresses piRNA and prevents stochastic silencing of the *gfp* transgene.

Most importantly, our theoretical model offered a new prediction: worms defective in neuronal sensation would be deficient in the inheritance of all RNAi responses, including exogenous, target-specific, dsRNA-induced gene silencing, even in the absence of temperature changes and regardless of stochastic silencing responses (**Figure 5D, E**). To test this prediction and assess the potency of RNAi inheritance in animals with impaired neuronal sensation, we exposed wild-type and *cmk-1(-)* animals – both carrying *mjIs134[mex-5::GFP]* germline GFP reporter transgene, which, unlike *bnIs1*, drives stable expression in germ cell nuclei at 20°C – to anti-*gfp* dsRNA and monitored the transgenerational silencing of GFP. We found that, while the basal expression *mex-5::GFP* remains unchanged in *cmk-1(-)* mutants (**Extended Figure 8A)**, these mutants are heritable RNAi deficient (“*Hrde*”), even at normal temperature (20°C) (**Figure 5F** and **Extended Figure 8B**). This, and our previous results, strongly demonstrate that neuronal perception of the environment, independent of the actual temperature or the biophysical effects of heat, could modify transgenerational inheritance of epigenetic information.

## Discussion

In this study, we found that perception of temperature via the AFD neurons affects germline small RNAs cell non-autonomously, influencing the dynamics of transgenerational epigenetic inheritance. AFD controls the expression of small RNA factors, including Argonaute proteins and RdRPs, as well as heritable RNAi through FLP-6 neuropeptide and serotoninergic signaling, which directly or indirectly affects the activity of transcription factor HSF-1 in the germline (**Extended Figure 9**).

In this manuscript, we used three complementary model systems to interfere with neuronal perception of temperature: a genetic knockout model, an ablation model, and a chemogenetic model. Moreover, our in-depth genetic analysis supports the conclusion that the thermosensory neural circuit, involving AFD neurons, CMK-1, and downstream neuromodulators, modulates germline Argonaute activity, thereby affecting heritable gene expression memory. In addition to AFD, it is likely that other sensory neurons are involved in promoting germline gene silencing. Supporting this notion, animals lacking *flp-6*, which is expressed in other amphid neurons outside of AFD, including ASE and ASG neurons^95^, showed a stronger a phenotype than AFD triple mutants. Similarly, knocking out *cmk-1*, which is expressed in multiple neuronal types^44^, causes a bistable expression pattern when the mutants were transferred from low to high temperature – a phenotype absent in most thermosensory mutants we analyzed in this work. While we cannot rule out the possibility that disrupting neuronal signaling may trigger a systemic stress response that ultimately impacts germline RNAi pathways, this does not contradict our model that regulatory signals originating from a single pair of AFD thermosensory neurons can generate perduring transgenerational effects.

This work supports our previous finding on the roles of HSF-1 in regulating heritable small RNA responses^75^. We showed that disrupting AFD thermosensation and eliminating HSF-1 relieved epigenetic silencing in the germline. Previous studies have demonstrated that HSF-1 has at least two roles – a developmental program and a stress-response program – both of which compete for HSF-1 binding^87,96,97^. While we did not focus on the molecular details of HSF-1 regulation in this manuscript, other works have indicated that HSF-1 binds the promoters of some epigenetic genes under non-heat shock conditions^87^. Consistently, we found that germline-specific depletion of HSF-1 causes downregulation of genes encoding Argonautes and other epigenetic regulators. Future studies are required to understand how HSF-1 and the small RNA machinery co-regulate development and stress responses.

In this work, we zeroed in on the perception of temperature due to the profound roles temperature play in development, reproduction, and aging^47,98–100^. However, other neuronally perceived stimuli – such as other types of stress, positive cues, or even neuronally-controlled learned behaviors – may be able to affect small RNA silencing, similar to the effects of temperature we reported here. Our approach – disrupting the function of specific neurons using complementary strategies and characterizing environment-sensitive transgenerational silencing of specific genes – offers a straightforward paradigm for studying neuron-to-germline communication in regulating inheritance. Future research could aim to decipher the neuronal ‘code’ that modulates epigenetic inheritance and characterize its physiological relevance.

It is likewise possible that different regimes of temperature stimuli, such as low temperature exposure (12-19°C), a transient heat shock, or recurring temperature fluctuations, may have different repercussions on small RNA inheritance. For example, our previous work has shown that a 2-hour heat shock is sufficient for “resetting” of ancestral small RNA responses^101^. In this work, we focus on the effects of prolonged (> 5 generations) exposure to elevated temperature, which leads to downregulation of epigenetic genes and de-repression of many genomic loci. This repression gradually recovers over several generations after animals are returned to lower temperature. Future studies may investigate how fluctuating environmental conditions shape the small RNA pool and influence transgenerational epigenetic inheritance.

Why would worms actively repress small RNA pathways when exposed to high temperatures? Reducing epigenetic regulation could constitute a mechanism for increasing genetic and phenotypic variability in largely isogenic population. *C. elegans* can distinguish between self and non-self genes, actively repressing foreign elements using heritable small RNAs^21,102^. In normal conditions, the activity of small RNA machinery is required for maintaining genome stability and preventing the harmful effects of transposition. When faced with stress, on the other hand, inhibition of the small RNA machinery may induce phenotypic plasticity and/or novel genetic variants, potentially allowing the animals to adapt to the environments^103^.

The nervous system of *C. elegans* in general, and the AFD neurons in particular, were shown to mediate a variety of cell-nonautonomous processes^45,46,104–107^. Persistent epigenetic memory of temperature stress in the absent of the initial trigger has been shown to have a negative fitness cost, at least under laboratory conditions^32^. It is tempting to speculate that intact thermosensation is required for re-calibration of the epigenetic states, allowing animals to readily adapt to new environments and promote the survival fitness of future generations – an intriguing model that warrants future investigation. In summary, the findings of this study illuminate a novel mechanism by which neuronal activity could control germline gene expression and direct non-Mendelian inheritance.

## Materials and Methods

### Worm cultivation

We used standard culture techniques to maintain the nematodes. We grew the worms on Nematode Growth Medium (NGM) plates and fed with OP50 bacteria. We took extreme care to avoid contamination or starvation for at least five generations prior to each experiment. We discarded contaminated or starved plates from the experiments. We performed all experiments with at least three biological replicates. We indicated all of the nematode strains we used in this study in the **Extended Table 5.**

### Histamine treatment

In experiments where worms were treated with histamine, we added a sterile-filtered 1M solution of histamine dihydrochloride to NGM agar at 50°C immediately before pouring the plates, to a final concentration of 10mM^67^. We stored histamine plates at 4°C for up to a month before usage.

In experiments where worms lost the HisCl1 extrachromosomal construct, we validated that all P-1 parents of worms lost the construct via PCR amplification of the HisCl1 construct, discarding any plate that showed residual presence of HisCl1.

### Fluorescence microscopy

For fluorescent microscopy assays, we used either an *Olympus IX83* motorized inverted wide-field microscope, or an *Olympus BX63* motorized upright wide-field microscope. We imaged experiments with a 10X objective lens, with an exposure time of 1000ms for GFP and 1500ms for mCherry.

Generally, we picked worms onto a microscope slide with 2% agarose pad containing drops of levamisole to induce paralysis, covered them with a glass coverslip, and imaged them after 2-5 minutes.

### Scoring of germline GFP reporter silencing

We scored GFP silencing and inheritance using a binary system: no visible expression (OFF), or any level of expression (ON). The results of each experimental group within each biological replicate were summarized as the percentage of worms showing any GFP expression, out of the entire worm population 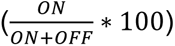. In a typical experiment, we performed at least three biological replication and at least two technical replicates. Each data point consists of at least 30 animals. The experimenters were blinded to the experimental conditions while scoring the experiments using DoubleBlind, a custom software tool that automatically and reversibly replaces file names with random strings (https://github.com/GuyTeichman/DoubleBlind).

In experiments and experimental conditions expressing HisCl1 constructs, we discarded worms that exhibited a non-specific expression pattern or no visible expression in the appropriate neuron pairs from the analysis.

In experiments where we quantified the basal expression level of GFP (**Extended Figure 11A**), we used the ImageJ Fiji ‘measure’ function^108^ to measure the integrated density of the three germline nuclei closest to spermatheca in each worm, as well as a mean background measurement of the worm in the germline’s vicinity. If less than three germline nuclei were visible, we took the remaining measurements in the estimated location and size of the germline nuclei instead. We calculated the corrected total cell fluorescence (CTCF) of each germline nucleus as previously described^101^. We used the mean of all three CTCF scores as the worm’s fluorescence score, and normalized the scores of each biological replicate to the mean of the control condition in that biological replicate.

Since some genetic background/experimental conditions can lead to developmental delays, worms in different conditions sometimes reached adulthood on different days. To avoid age bias between the different conditions, we tracked the developmental stages of all worms, and imaged them when they reached the ‘day-one adulthood’ stage.

### Statistical analysis

For transgenerational silencing experiments, we analyzed the data using a mixed effect model, with the genotype/treatment considered as a categorical factor, and the generation considered as repeated-measures data. We then followed up with multiple comparisons, comparing the means of the experimental conditions to the mean of the control condition within each generation.

In experiments with exactly two conditions (experimental vs. control), we used the Unpaired Student’s t-test with Welch’s correction for unequal variances. For experiments with multiple conditions and/or generations, we corrected for multiple comparisons using the Two-stage step-up method of Benjamini, Krieger and Yekutieli^109^.

### mRNA and small RNA sequencing experiments

#### Collecting worms for sequencing

For the *gcy-8/18/23(-)* mutant experiment, we raised wild-type and mutant animals in 25°C for at least 5 generations. We ran the experiment in three biological replicates, each containing 3-5 technical replicates. We initiated each technical replicate with 25 parent worms in the P-1 generations. After 2-3 hours of synchronized egg laying, we imaged the parent worms to ensure a uniform starting point of GFP reporter expression. We additionally imaged the worms in every generation, to keep track of the GFP expression phenotype of each technical replicate. We synchronized each new generations from 25 randomly selected parent worms. We isolated adult worms in the P0, F1, F2, and F3 generations.

From each biological replicate, we selected one wild-type and one mutant replicate, based on their GFP reporter expression in the F3 generation: we selected the replicate whose GFP expression was closest to its genotype’s median GFP expression. We did this to ensure that the samples we chose were representative of the phenotype we observed in previous experiment, while avoiding selection bias.

For the *cmk-1(-)* mutant experiment, we raised wild-type and mutant animals at 20°C for at least 5 generations. We ran the experiment in three biological replicates, each containing 8-10 technical replicates. We initiated each technical replicate with 25 parent worms in the P-1 generations. After 2-3 hours of synchronized egg laying, we imaged the parent worms to ensure a uniform starting point of GFP reporter expression. We additionally imaged the worms in the F2 generation, to keep track of the GFP expression phenotype of each technical replicate. We synchronized each new generations from 25 randomly selected parent worms. We isolated adult worms in the P0 and F3 generations.

From each biological replicate, we selected two technical replicates to be sequenced based on their GFP expression in the F2 generations: we selected the worms with the highest GFP expression (“expressing” samples), and the worms with the lowest GFP expression (“silencing” samples). We did this to ensure that we picked worms representing both extremes of the silencing phenotypes we observed in previous experiments and facilitate comparisons between worms that responded differently to changes in temperature.

### RNA extraction

We performed RNA extraction as previously described^54^. We lysed worms using the TRIzol reagent. We added 400 µl of TRIzol to 100 µl of adult worms and performed three cycles of freezing in -80°C and vortexing at RT for 15 minutes. We added 100 µl of chloroform to the samples, transferred them to pre-spun Heavy Phase-Lock tubes, and rested them on ice for 10 minutes. Next, we centrifuged the tubes at 16,000 g for 15 minutes at 4°C. We transferred the aqueous phase to a second pre-spun Heavy Phase Lock tube, added 1:1 of a Phenol:Chloroform:Isoamyl Alcohol (25:24:1) mixture, and centrifuged the samples at 16,000 g for 5 minutes at RT. We transferred the aqueous phase to a 1.5 ml Eppendorf tube and added 20 µg of Glycogen and 1:1 Isopropanol. We incubated the samples at -20°C overnight, and then spun them for 30 minutes at 16,000 g at 4°C. We washed the resulting pellet twice with ice-cold 70 % ethanol and then air-dried for 10 minutes. We then re-suspended the pellet in 20 µl of RNase-free water. We validated the concentration of the total RNA using Qubit RNA high-sensitivity kit, and ensured that the RNA integrity was sufficiently high (RIN >= 7) using an Agilent 4150 BioAnalyzer instrument and High Sensitivity RNA ScreenTapes.

### mRNA libraries

We enriched the samples for poly-A-tailed mRNA molecules using the NEBNext Poly(A) mRNA Magnetic Isolation Module, using a starting quantity of 100-1000 ng Total RNA.

For the *cmk-1* mRNA samples, we generated cDNA libraries using the NEBNext Ultra II Directional RNA Library Prep Kit and the NEBNext Dual Index Primers multiplex oligos. We sequenced the libraries using an Illumina NextSeq 500 instrument.

### Small RNA libraries

We treated total RNA samples with RNA 5’ Polyphosphatase (LGC, Biosearch Technologies) to ensure 5’ monophosphate-independent capture of small RNAs. We prepared small RNA libraries using the NEBNext® Small RNA Library Prep Set for Illumina® according to the manufacturer’s protocol. We separated the cDNA libraries on a 4 % agarose E-Gel (Invitrogen, Life Technologies), and selected the 140-160 nt length bands. We pooled and purified the size-selected cDNA using the MinElute Gel Extraction kit (QIAGEN). We sequenced the libraries using an Illumina NextSeq 500 instrument.

### RNA sequencing analyses

Sequencing analyses were done using RNAlysis^110^, and all of the details of the analyses, including workflow, function parameters, graphs, and intermediate data, are available in **Extended Files 2-6 – RNA Sequencing Interactive HTML Report**.

For small RNA sequencing datasets, we used FastQC to assess sample quality, filtered out reads shorter than 15 nucleotides after adapter trimming using CutAdapt^111^, aligned small RNA reads using ShortStack^112^, and counted aligned small RNA reads using FeatureCounts^113^. Differential expression analysis was performed using Limma-Voom.

For mRNA sequencing datasets, we used FastQC to assess sample quality, aligned reads using HISAT2^114^, and counted aligned reads using FeatureCounts. Differential expression analysis was performed using DESeq2^115^.

### Classification of groups of siRNAs that accumulate transgenerationally at 20°C

First, to identify groups of siRNAs whose expression changes significantly over generations at 20°C, we ran differential expression analysis using Limma-Voom, with the design formula “∼ Genotype + poly(Generation, degree = 2)”. We kept only siRNAs whose polynomial coefficients of the *Generation* parameter were significantly different from 0 (q < 0.1).

Next, to group these siRNAs based on their change pattern, we filtered genes based on the values of their 1^st^ and 2^nd^ polynomial coefficients, keeping only those whose 1^st^ and 2^nd^ coefficients were larger than 0.5 or smaller than -0.5. We then examined the four possible combinations of coefficient ranges (1^st^/2^nd^, greater than 0.5/lesser than -0.5), and picked the two groups of siRNAs that exhibited transgenerational accumulation dynamics.

### Mathematical model

We based our mathematical model on the Toggle-Inhibitor-Competition (TIC) model developed by Karin et al.^93^, which aimed to explain transgenerational inheritance of small RNA responses, as well as the similarities and differences in small RNA inheritance between isogenic worms. We expanded this model to apply to temperature-dependent small RNA-mediated gene silencing and simulated the effects of the changes in the model’s parameters and components affecting this process.

Based on our experimental results, we aimed to fulfill the following requirements:

1. Change of growth temperatures modifies the small RNA responses of worms in both sensation-dependent and -independent manners.
2. The *gfp* reporter should acquire silencing gradually and stochastically when transitioned from high to low temperatures, but de-silence rapidly and uniformly when transitioned from low to high temperatures.
3. Worms defective in AFD temperature sensation show delayed silencing at low temperatures, but de-silence normally at high temperatures.

Based on our mRNA and small RNA sequencing results, we made the following assumptions:

1. High temperatures, or defective temperature sensation, reduce the silencing capacity of worms (**Vtot**) by ∼two-fold. We base this on the observed downregulation of small RNA machinery genes in 25°C, and in worms defective in AFD temperature sensation (**Figure 3**). This is also supported by **Figure 3A** of, which demonstrates that RNAi inheritance is reduced when worms are raised in high temperatures in the generation following RNAi.
2. To explain the RNAi-free accumulation of silencing in low temperatures, the *gfp* reporter gene must be targeted by stochastic silencing events at a higher frequency than other genes. According to our small RNA sequencing data, it is in the top 1% of small RNA-targeted genes – normalized siRNA levels are ∼20-25 fold larger than that of the average protein-coding gene (**Extended Table 6**).
3. To explain the maintenance of near-100 % expression of the *gfp* reporter at high temperatures, the frequency of *gfp* silencing events must be reduced by high temperatures. This is possibly due to reduction in piRNA levels (and probably other factors), which we observed in our small RNA sequencing data (**Figure 4C**).

We then proceeded to simulate transgenerational-silencing experiments, where worms are transitioned from a higher temperature to a lower temperature, or vice versa. As in the “real” experiments, we started each simulated experiment with 25 random worm lineages, and randomly picked 25 worms at each generation (84 hours) to spawn the next generation of worms.

For the simulated RNAi experiment, we initiated an RNAi trigger as in Karin et al. 2023, with the entire simulation being conducted in a lower temperature, such that the silencing capacity (Vtot) of wild-type worms is ∼2 times higher than in the mutants.

The simulation code is available online at github.com/GuyTeichman/Teichman_2024, as well as **Extended File 1 – simulation code**.

### DNA constructs and transgenic animals

To express HisCl1 and GCaMP6 in the AFD neurons we used the AFD-specific promoter *gcy-8*^60^. An expression vector containing *gcy-8p::HisCl1::SL2:mCherry:unc-54 3’UTR* was generously gifted by the lab of Dr. Piali Sengupta. An expression vector containing *gcy-8p::GCaMP6:unc-54 3’UTR* was generously gifted by the lab of Dr. Daniel Colón-Ramos.

To improve the expression specificity of the construct, we replaced the *unc-54* 3’UTR sequence in both vectors with the *rab-3* 3’UTR sequence (656bp amplified from genomic DNA) via Gibson Assembly.

To express HisCl1 and GCaMP6 in the ASK neurons we used the ASK-specific promoter *sra-7*^68^. We amplified 4kbp upstream of the *sra-7* ORF from *C. elegans* genomic DNA using the primers ATAGCGGCCGCGAGAAATATTTGATGGATGTTTG (forward) and ATTGGGATCCCAAAAGTCAACGGACTGTGA (reverse), which also inserted the restriction sites for NotI and BamHI respectively. We used Phusion® High-Fidelity DNA Polymerase (NEB) to amplify the *sra-7* promoter using the company’s protocol. We purified the resulting PCR product (QIAquick PCR Purification Kit, Qiagen) and double-digested it using NotI-HF and BamHI-HF (rCutSmart buffer, NEB) following the manufacturer’s standard procedure. We then purified the inserts (QIAquick PCR Purification Kit, Qiagen).

We cut each of the vectors in a similar manner using the same restriction enzymes. We separated the resulting products in agarose gel (1 %) and purified them (Wizard SV Gel and PCR Clean-Up System, Promega). We then ligated the inserts and vectors with Quick Ligase (NEB) following the company’s standard procedure. We transformed the ligated constructs to TOP10 competent cells under selection using carbenicillin at 100 ug/ml (Sigma). After transformation, we extracted the plasmids (QIAprep Spin Miniprep Kit, Qiagen). We verified the plasmid sequences using Sanger sequencing with an M13 primer (AGCGGATAACAATTTCACACAGGA).

We injected the HisCl1 and GCaMP6 constructs into N2 worms, along with the co-injection marker *rol-6(su1006)* (pRF4) and DNA ladder (1kb DNA Ladder N3232, NEB), generating the three AFD-specific independent lines BFF95, BFF96, BFF97: *pigEx17*/*18*/*19*[*gcy-8p::HisCl1::SL2::mCherry::rab-3* 3’UTR + *rol-6p::rol-6(su1006)::rol-6* 3’UTR + *gcy-8p::GCaMP6::rab-3* 3’UTR], as well as ASK-specific line BFF126: *pigEx21*[*sra-7p::HisCl1::SL2::mCherry::rab-3* 3’UTR + *rol-6p::rol-6(su1006)::rol-6* 3’UTR + *sra-7p::GCaMP6::rab-3* 3’UTR].

We injected the constructs in the following concentrations: HisCl1 (25 ng/μl), GCaMP6 (25 ng/μl), pRF4 (25 ng/μl) and DNA ladder (25 ng/μl). We then crossed the various HisCl1 lines into worm strain SS747, which contains a GFP temperature-dependent silencing reporter.

### Bacterial strains and growth

For infection survival assays, a GFP-expressing derivative of the pathogen Pseudomonas aeruginosa strain PA14 was used^116^. PA14-GFP bacteria were grown overnight at 37 °C in King’s B medium supplemented with 100 µg/mL rifampicin without agitation. 10 µL of this culture was then spread uniformly on 35 mm Slow Killing Plates (SKPs)^117^ using a bent glass transfer pipette, and incubated overnight at 37 °C. Plates were allowed to reach room temperature before transferring worms for infection.

### Temperature shift experiments and survival assays on PA14

Experimental groups included worms that were continuously maintained at 15 °C or 25 °C on OP50 for at least five generations. In temperature shift experiments, P0 mothers raised in designated temperatures were shifted at the L4 stage to a new temperature (from 15 °C to 25 °C, or from 25 °C to 15 °C) to ensure F1 egg development at the new temperature. P0 gravid worms were transferred to new OP50 plates and allowed to lay eggs for one hour before removal, to allow tight cohort synchronization of F1 progeny. F1 worms were maintained at the new temperature until the L4 stage, then transferred to PA14 SKPs for survival analysis. For F2 shifted animals, the progeny of the F1 generation were also maintained at the new temperature until the L4 stage and transferred to PA14 SKPs. Timed egg lays were staggered at the different temperatures so that all worms reached the L4 stage simultaneously for concurrent survival analysis.

L4 worms transferred for PA14 survival assays were briefly washed in M9 buffer and approximately 100 individuals were transferred by sterile glass pipette, split between three PA14-seeded SKPs. Infections were carried out at 25 °C for the duration of the assay, regardless of prior rearing temperature. Live worms were counted 1-2 times per day. Worms desiccated on the side of plates were censored from analysis. Statistical significance was assessed by a log-rank test after Kaplan-Meier survival analysis, performed in R.

## Acknowledgements

We thank all members of the Rechavi lab and co-authors for fruitful discussions, feedback, and support. We are especially grateful to Mélanie Bazin-Gélis and Thi Kim Thanh Vuong-Brender from the Schumacher lab for their help in generating strain BJS1456 and for their valuable feedback. O.R. is grateful to funding from the Eric and Wendy Schmidt Fund for Strategic Innovation (Polymath Award #0140001000) and the generous support from the Morris Kahn foundation. GT is grateful to the Milner Foundation. C.K.E was supported by EMBO Post-doctoral Fellowship (ATLF 6-2022). The Rechavi lab is funded by ERC grant #335624 and the Israel Science Foundation (grant#1339/17). O.R. and B.S. acknowledge funding from the John Templeton Foundation Grant (61734) and the Deutsche Forschungsgemeinschaft (project 437407415, SCHU 2494/10-1).

## Author contributions

O.R conceptualized the idea for the project. G.T., M.S., C.K.E., and O.R. conceptualized and designed the experiments. G.T., C.K.E., M.S., H.G., G.A, H.D., S.A., V.P., M.MG. and M.S. conducted the experiments. I.R. and O.S. generated the DNA constructs. V.P. and M.O.S. conducted calcium imaging experiments. G.T. and C.K.E. worked on Statistical Analysis, Data Curation, and Visualization. G.T. and H.G. ran bioinformatic analyses. G.T. modified the mathematical model and wrote the simulations. G.T. and C.K.E. wrote the original draft of the manuscript. G.T., C.K.E, H.G., P.R.S., D.H.M, B.S., and O.R. review and edit the manuscript.

## Data and Code Availability

A complete report of RNA sequencing data analysis, including intermediate files and function parameters, is available as **Extended Files 2-6 – interactive analysis reports**.

Small RNA and mRNA sequencing libraries generated during this study are available at the GEO repository accession number GEO: [TBA]

We used the following previously published datasets:

Apfeld J (2022) NCBI BioProject ID PRJNA822361. Neuronal temperature perception induces specific defenses that enable *C. elegans* to cope with the enhanced reactivity of hydrogen peroxide at high temperature.

Li J (2021) NCBI BioProject ID PRJNA680496. Using Auxin-inducible-degradation System to Interrogate Tissue-specific Transcriptional Programs of HSF-1 in Reproduction and Heat Shock Response.

## Extended data figures

**Extended Figure 1:**
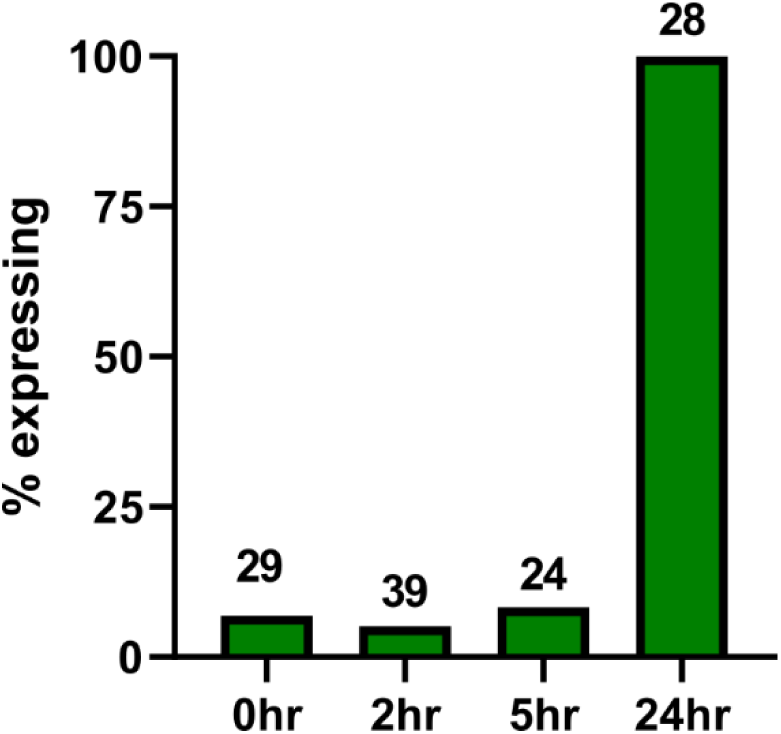
Heat-induced expression of *bnIs1* transgene. *bnIs1* is de-repressed when adult animals were grown at 25°C for at least 24 hours. Number of animals scored at each time point is indicated.

**Extended Figure 2:**
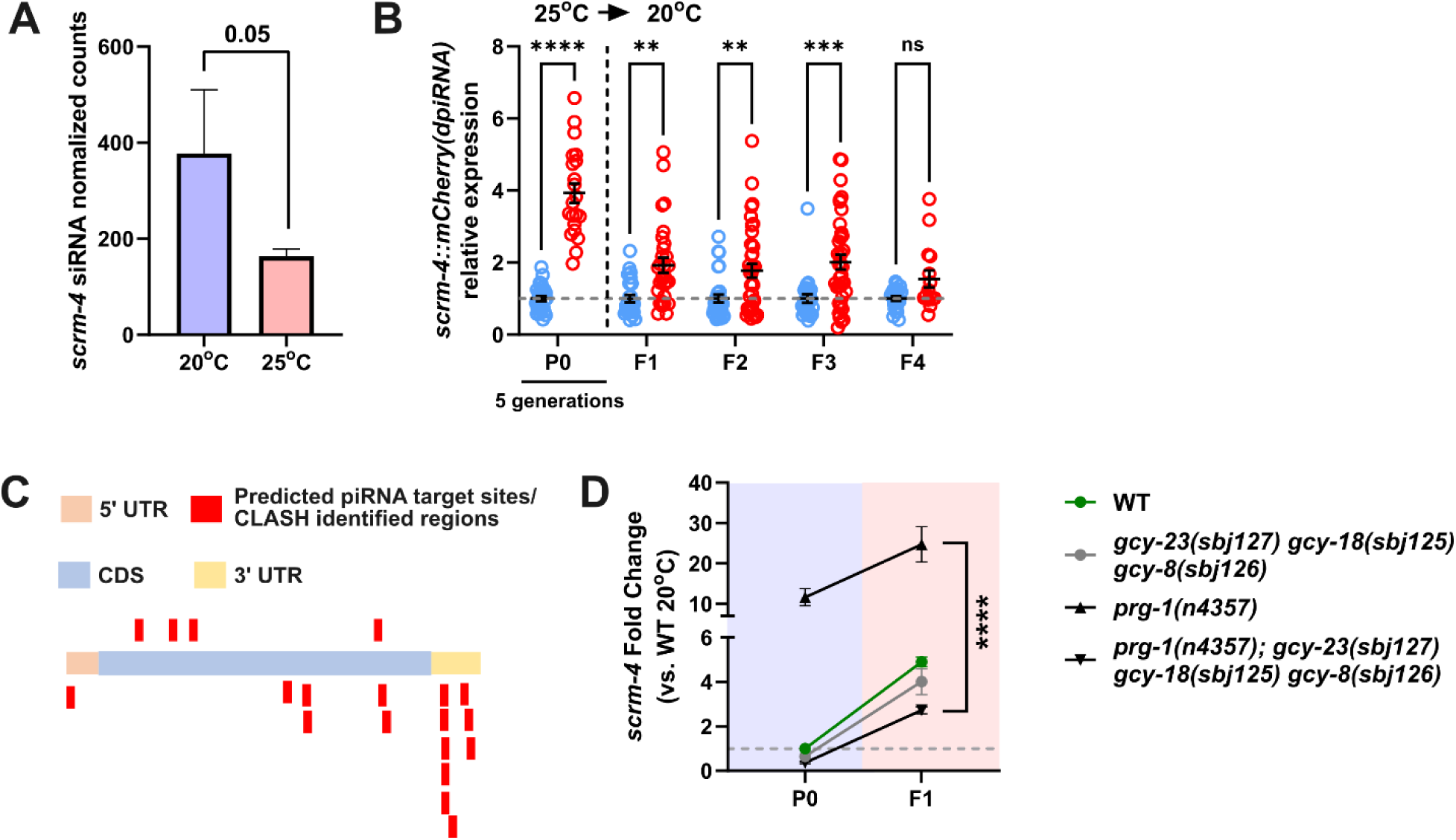
*scrm-4* serves a reliable RNAi sensor. (A) *scrm-4* siRNA is downregulated at 25°C compared to 20°C (B) *scrm-4* shows transgenerational silencing upon transitioning from 25°C compared to 20°C. (C) piRNA targeting sites in *scrm-4*, based on piRTarBase. (D) Knocking out AFD triple gcy rescues overexpression of *scrm-4* in *prg-1(-)*. (mean ± SEM). (*) indicates q < 0.05, (**) indicates q < 0.01, (***) indicates q < 0.001, and (****) indicates q < 0.0001

**Extended Figure 3:**
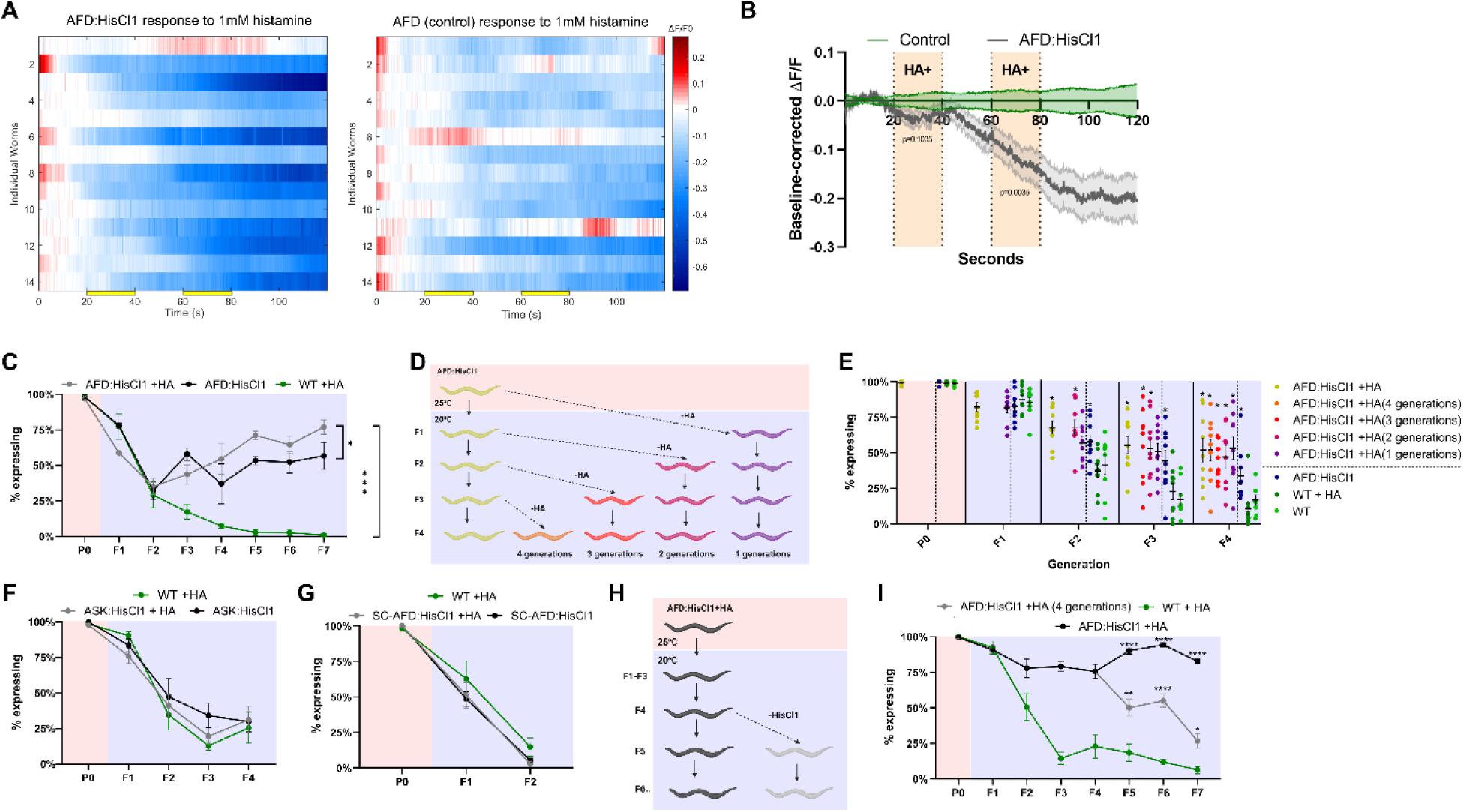
HisCl1-mediated hyperpolarization affects temperature-dependent transgene silencing in an AFD-specific manner. (A, B) Histamine exposure reversibly hyperpolarizes HisCl1-expressing AFD neurons. Activity of the AFD sensory neurons was quantified by GCaMP6s fluorescence intensity in a microfluidic device controlling histamine exposure. Wild-type (n=14) and AFD:HisCl1 (n=14) worms were loaded into chips and exposed to 1mM HA for two periods of 20 seconds (20-40 seconds and 60-80 seconds, see yellow bars in A). (C) Hyperpolarization of the AFD neurons suppresses transgene silencing upon transition to a low temperature, with the response lasting for at least 7 generations. (D) Experimental scheme. AFD:HisCl1 and control worms were transitioned from a high to a low temperature. After each generation in the low temperature, some worms from the AFD:HisCl1+HA group were transitioned to plates no longer containing histamine, and were then tracked for more generations in those conditions. This led to the creation of five experimental groups, that were exposed to histamine for 5/4/3/2/1 generations in total. In the purple group, in which the worms experienced only one generation of histamine treatment, delayed silencing was evident especially in F4. Significance was assessed relative to AFD:HisCl1. Additional comparisons were made between AFD:HisCl1 and wildtype. (E) Histamine treatment of AFD:HisCl1 worms suppresses temperature-dependent silencing, even when the histamine treatment is stopped for 1-3 generations. (F) Expression of HisCl1 and hyperpolarization of the ASK sensory neurons do not affect temperature-dependent transgene silencing. (G) Single-copy expression of the histamine-gated chlorine channel HisCl1 under the *gcy-8* promoter does not suppress transgene silencing upon transition to a low temperature. (H) Experimental scheme. AFD:HisCl1 and control worms were transitioned from a high to a low temperature. After 4 generations in the low temperature, some worms from the AFD:HisCl1 group were selected to lose the extrachromosomal AFD:HisCl1 construct and were then tracked for three more generations in a low temperature. (I) Worms whose ancestors expressed AFD:HisCl1 maintain GFP expression for at least 3 generations after losing the AFD:HisCl1 construct. In all relevant panels, error bars indicate mean ± SEM. (*) indicates q < 0.05, (**) indicates q < 0.01, (***) indicates q < 0.001, and (****) indicates q < 0.0001

**Extended Figure 4:**
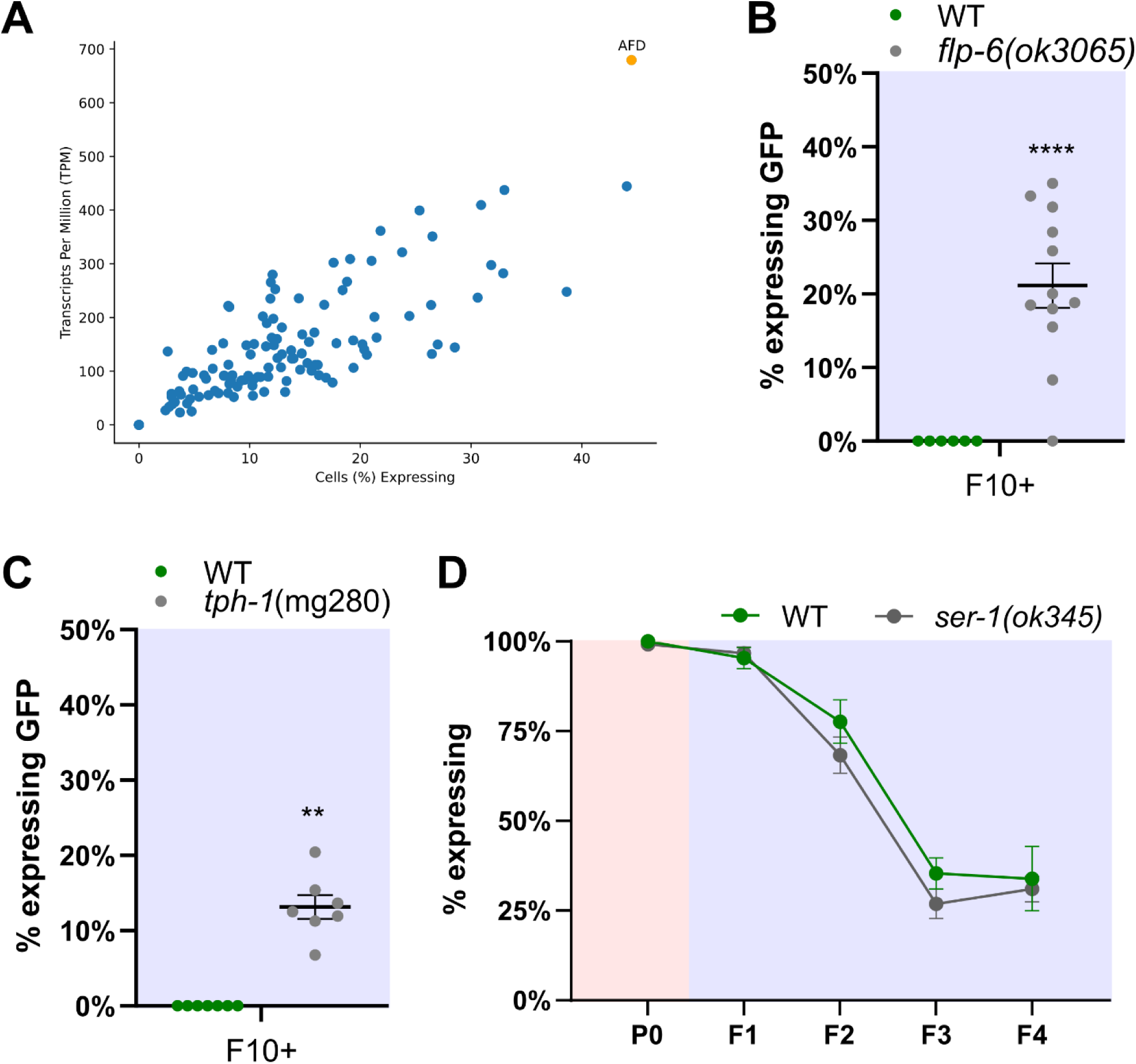
The long-term effects of the AFD thermosensory circuit on small RNA silencing. (A) The CaM Kinase gene *cmk-1* is expressed chiefly in the AFD neurons. Data is taken from the CeNGEN Project. Shown is a scatter plot comparing the average TPM of *cmk-1* in various neuron classes (Y-axis) to the percentage of cells of various neuron classes with detectable expression of *cmk-1* (X-axis). Each dot represents a neuron class, with the orange dot representing the AFD neuron class. (B, C) Knocking out the neuropeptide gene *flp-6* and the tryptophan hydroxylase gene *tph-1* inhibits temperature-dependent small RNA-mediated gene silencing for at least 10 generations following transition from high to low temperature. (D) Knocking out *ser-1* does not affect silencing of *bnIs1*. In all relevant panels, error bars indicate mean ± SEM. In (B) and (C), each dot represents a replicate of the experiment that consists of at least 30 animals. (**) indicates q < 0.01 and (****) indicates q < 0.0001.

**Extended Figure 5:**
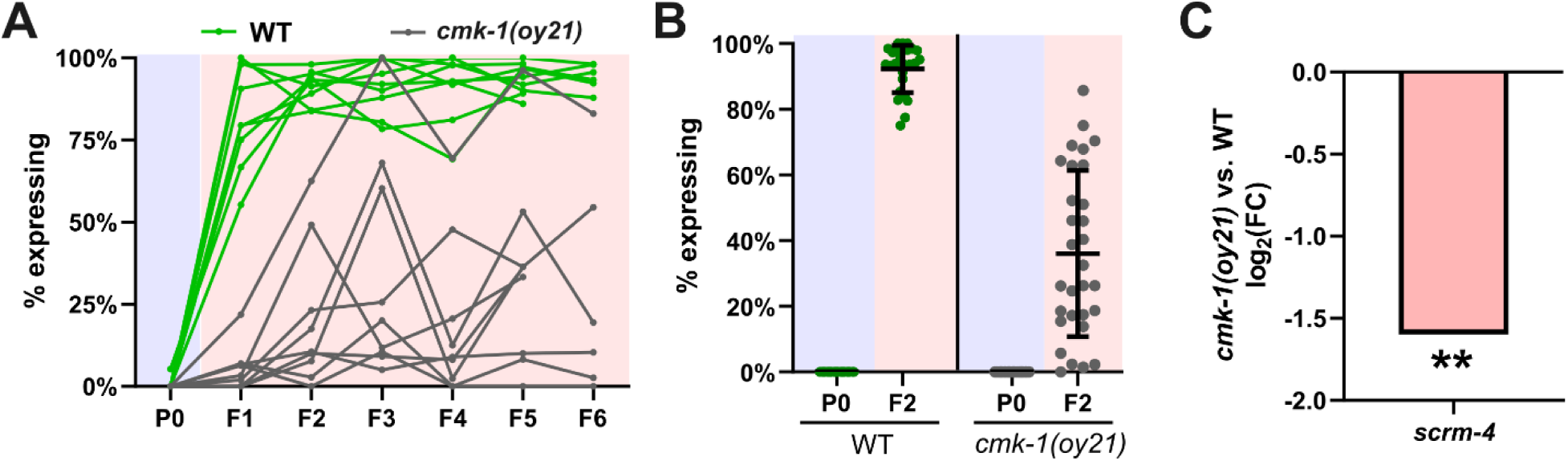
Mutants of the CaM Kinase gene *cmk-1* exhibit a high-variance de-silencing response in high temperatures. (A) Knocking out *cmk-1* causes high-variance de-silencing response when transitioning from 20°C to 25°C. Each line represents a replicate of the experiment, consisting of at least 30 worms per generation. See Figure 2B for summary data. (B) When transitioning to high temperatures, *cmk-1(-)* mutants exhibit a high-variance de-silencing response, with some lineages completely de-silencing the GFP transgene, while others maintain complete silencing. Out of 27 replicates, we picked the three most extreme samples in each direction for RNA sequencing (see **Materials and Methods** for full experimental protocol). Error bars indicate mean ± SEM. (C) *cmk-1(-)* mutants shows downregulation of *scrm-4*.

**Extended Figure 6:**
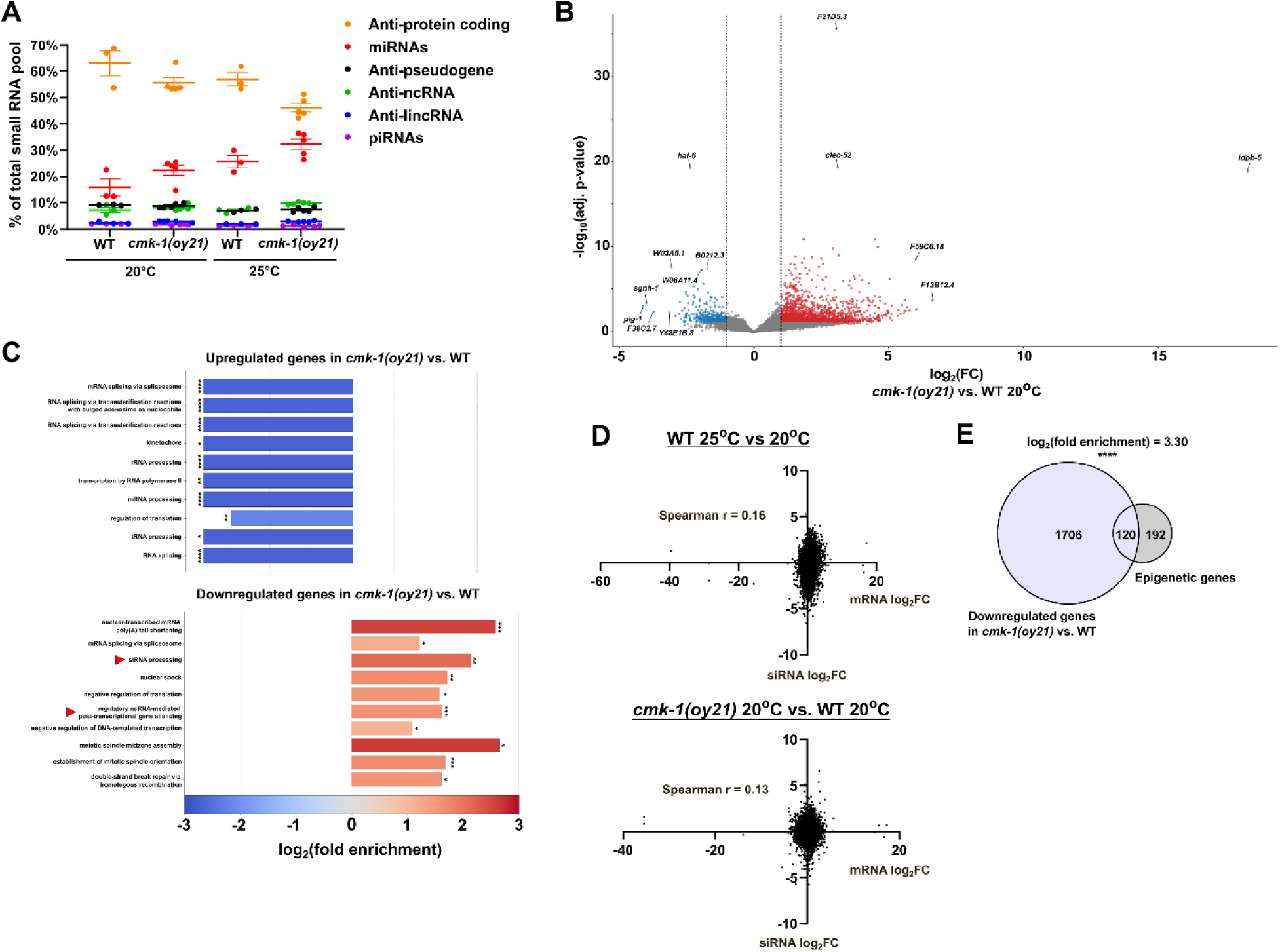
Small RNA and mRNA expression in *cmk-1(-)* mutants. (A) Growing at 25°C and mutation in *cmk-1* downregulates small RNAs targeting protein-coding genes. Shown are the percentage of the total small RNA pool aligned to different types of genomic features (mean ± SEM). Each dot depicts a biological replicate. (B) Volcano plot showing gene expression change in *cmk-1(-)* vs. WT at 20°C. (C) GO enrichment analysis of genes differentially expressed in *cmk-1(-)* vs. WT. Categories involved in small RNA-mediated gene silencing are highlighted. (D) Correlation between mRNA and siRNA expression across temperatures and between *cmk-1(-)* and WT. (E) Epigenetic genes tend to be downregulated in *cmk-1(-)* mutants.

**Extended Figure 7:**
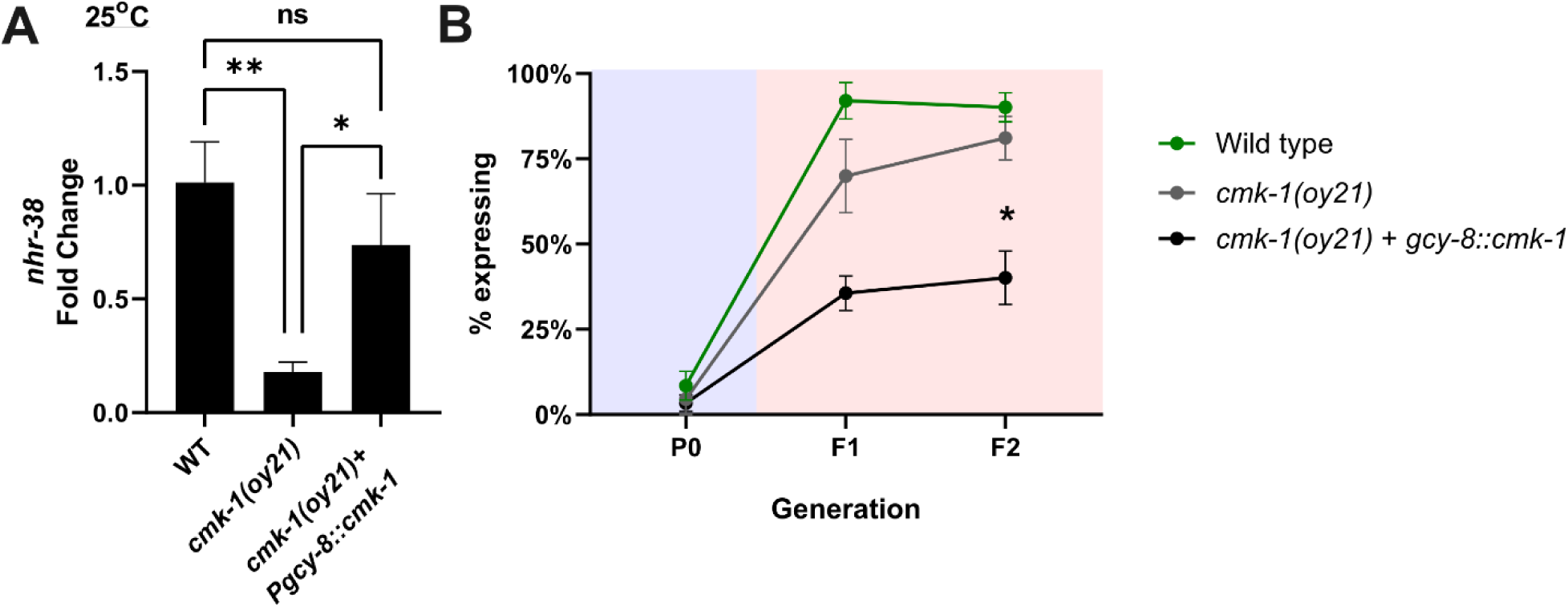
Rescuing *cmk-1* in AFD neurons causes enhanced silencing. (A) Expressing *cmk-1* under the control of *gcy-8* promotor rescues *nhr-38* expression in *cmk-1(-)*, indicating that the single-copy transgene is functional. (B) *cmk-1(-) + gcy-8::cmk-1* shows enhanced transgene silencing compared to wild type and *cmk-1(-)*. In all panels, error bars indicate mean ± SEM. (*) indicates q < 0.05, (**) indicates q < 0.01, (***) indicates q < 0.001, and (****) indicates q < 0.0001

**Extended Figure 8:**
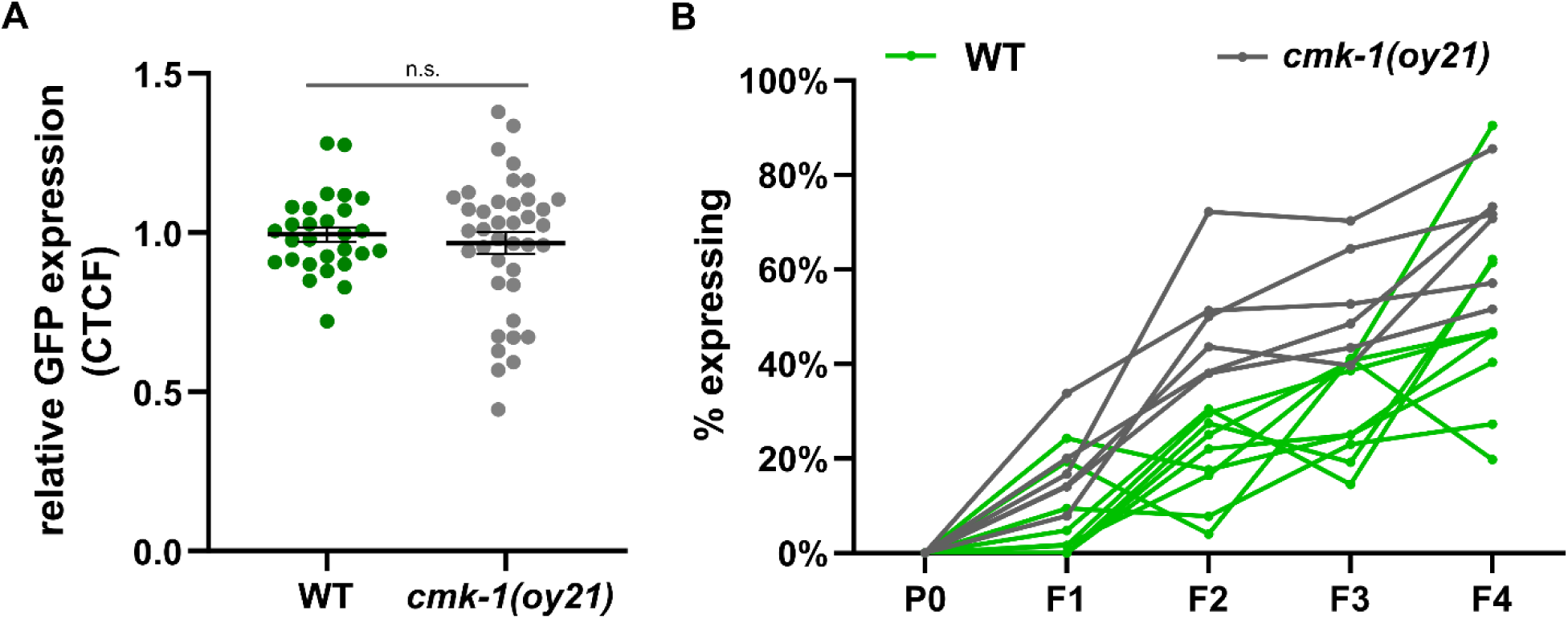
Modified temperature perception, even in the absence of temperature change, repressed heritable RNAi responses (individual trajectories) (A) The basal expression level of the *mjIs134*[*mex-5p::gfp::h2b::tbb-2*] transgene does not differ between wild-type worms and *cmk-1(-)* mutants. Error bars indicate mean ± SEM. Each dot represents the average of three measurements of an individual worm. (B) Defective sensory perception is sufficient to dampen transgenerational inheritance of RNAi responses. Each line represents a replicate of the experiment, consisting of at least 30 worms per generation.

